# Single-cell transcriptomics reveals cell-type-specific circadian rhythms and their disruption by acute misalignment in mouse aorta

**DOI:** 10.64898/2026.09.03.749170

**Authors:** Benjamin J. Auerbach, Ronan Lordan, Soon Y. Tang, Elizabeth Hennessy, Seán T. Anderson, Ujjalkumar Subhash Das, Ryan McConnell, Mingyao Li, Garret A. FitzGerald

## Abstract

The circadian molecular clock is a 24-hour cellular timekeeper that influences many features of cardiovascular function. Disruption of the circadian clock via misalignment with the light-dark cycle is associated with a higher incidence of cardiovascular disease and raises cardiovascular risk factors in humans. Nonetheless, the cell-type-specific molecular basis for how misalignment affects the vasculature remains poorly understood. To address this, we performed single-cell RNA-sequencing from whole mouse aorta at ZT0, ZT6, ZT12, and ZT18 under aligned and acutely misaligned (6-hour phase advance) light-dark cycles in both male and female mice. Leveraging Bayesian variational inference, we estimated posterior waveforms for 141,752 cells across four major cell types and identified hundreds of cycling genes in vascular smooth muscle cells (SMCs) and fibroblasts. Pathway and transcription factor enrichment analyses revealed coordinated circadian activity in cholesterol biosynthesis, smooth muscle contraction, and extracellular matrix organization. Notably, SMC genes implicated in phenotypic switching showed coordinated temporal patterns, with genes promoting switching peaking at dusk and genes restraining switching peaking at dawn. Comparing males and females, we found that female SMCs are broadly more rhythmic, with higher amplitudes and nearly twice as many cycling genes after controlling for cell counts and library sizes—a sex difference that was cell-type-specific and not observed in fibroblasts. After acute misalignment, cycling genes showed reduced amplitudes, and peak times showed limited adaptation to the new light-dark cycle. Given that the central clock is known to adapt near-completely during this timeframe, these observations suggest internal misalignment between central and peripheral rhythms. Moreover, altered relative timing of core clock genes within cells indicates that misalignment is created at the intracellular level as well. In SMCs, gene expression patterns were consistent with proteostatic stress, including broad downregulation of protein chaperones and stress-response genes, which cells appear to cope with by upregulating protein degradation pathways. In parallel, in vivo vascular phenotyping showed increased vascular permeability in both sexes, reduced urinary nitrate in males, and increased microvascular thrombus formation in males following acute misalignment. This atlas provides a resource for understanding how circadian misalignment disrupts vascular homeostasis and may contribute to cardiovascular disease risk.

## Introduction

The circadian molecular clock is a 24-hour timekeeping mechanism found in nearly every cell in humans. The time of the clock, referred to as circadian phase, is determined by the mRNA and protein concentrations of the clock’s constituent genes, referred to as clock or core clock genes. Clock genes are organized in a transcriptional-translational feedback loop that enables cells to maintain self-sustained *∼*24-hour oscillations in the concentrations of clock gene mRNA. Clock gene proteins additionally interact with cell-type-specific regulatory factors to drive rhythmic transcription of genes referred to as clock-controlled genes (CCGs). It is largely through these CCGs that circadian clocks generate rhythmic cellular and organismal behaviors.

Many features of cardiovascular function are under direct control of the circadian clock, such as vascular tone [1], thrombus formation [2], endothelial nitric oxide synthase activity [3], and myeloid cell recruitment and adhesion [4, 5]. Crucially, disruption of circadian timing contributes to vascular dysfunction and increases cardiovascular disease risk. Chronic disruption of the clock via light-dark (LD) misalignment by forced cycle desynchrony protocols increases cardiovascular risk factors in humans [6] and epidemiological studies associate shift work with an increased incidence and prevalence of cardiovascular disease [7, 8, 9]. However, these chronic outcomes arise from repeated episodes of acute circadian misalignment, during which environmental and behavioral cycles become uncoupled from the endogenous circadian clock. The cell-type-specific transcriptional responses that occur during this early period of adaptation remain poorly understood. In humans, biomarkers of cardiovascular risk differ between males and females. These include arterial stiffness [14], lipid profiles [15, 16], blood pressure [17], and immune phenotypes [18]. Features of vascular disease also differ between sexes, such as in atherosclerotic plaque size, composition, and stability [19, 20]. Existing literature suggests females may exhibit more robust circadian characteristics than males and their clocks appear more resilient to disruption [21]. Considering this, and given the clock’s influence over key vascular disease risk factors [22, 23, 24], characterizing how the sexes differ in their molecular response to clock perturbations will be important to understand whether sex differences in vascular disease presentation are mediated by the circadian clock.

In this study, we combined circadian single-cell RNA sequencing and in vivo vascular phenotyping to determine how acute circadian misalignment reshapes cell-type-specific transcriptional rhythms in the mouse aorta. We established a circadian atlas of aortic cell types, uncovered sex-dependent differences in rhythmic gene expression, showed that acute phase advance disrupts circadian transcriptional programs, which may be linked to impaired vascular function.

## Results

### Generation of a single-cell mouse aorta circadian atlas

To investigate these questions, we performed single-cell RNA-sequencing (scRNA-seq) experiments from whole aorta samples harvested over circadian time courses (Figure 1a). First, to assess the contributions of sex and cell type on circadian expression of cells aligned with the light-dark (LD) cycle, whole aorta samples were collected from 12-week-old male and female C57BL/6 mice entrained to a 12:12 LD cycle for 2 weeks. Mice were sacrificed at Zeitgeber time (ZT) 0, 6, 12, or 18 hours, and whole aorta samples were harvested (Figure 1b). We additionally collected data from inducible Bmal1 global deletion models (Methods). To assess the response to acute circadian misalignment, 12-week-old male and female C57BL/6 mice were entrained to a 12:12 LD cycle for 2 weeks, after which the start of the LD cycle was advanced 6 hours (Figure 1c). Mice were sacrificed 5 days after the phase advance at ZT 0, 6, 12, or 18 hours. For all conditions, cells were dissociated into single-cell suspensions and profiled using the 10X Genomics Chromium platform.

**Figure 1:**
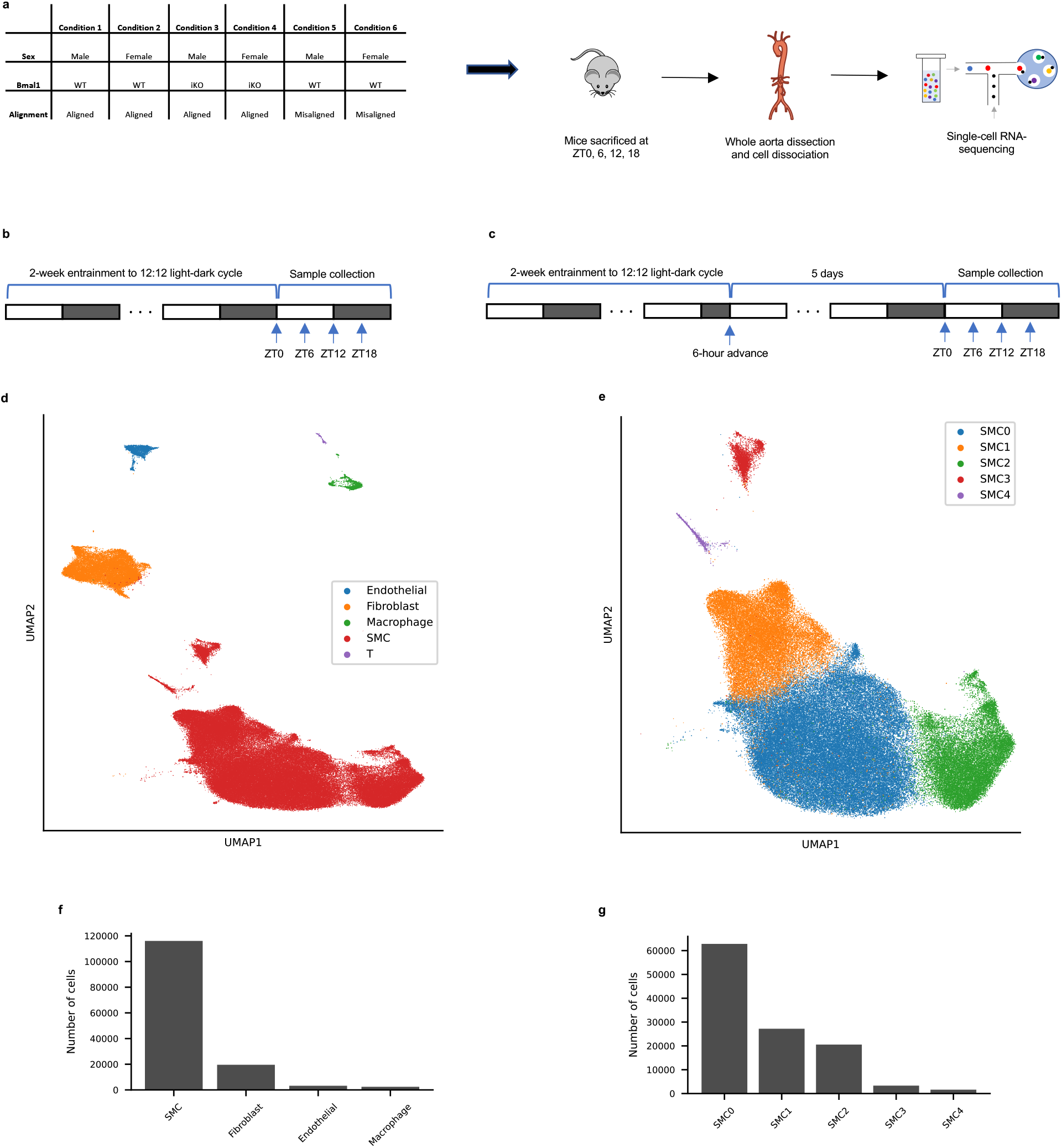
Experimental overview and cell types of the mouse aorta circadian atlas. a) Experimental conditions: male and female C57BL/6 mice under aligned and misaligned light-dark cycles, and male and female Bmal1fl/fl:CAGGcreERt2/+ mice under aligned conditions. Mice were sacrificed at ZT0, ZT6, ZT12, and ZT18, whole aortas were dissected and dissociated, and single-cell RNA-sequencing was performed. b) Light-dark cycle and sample collection protocol for aligned mice. c) Light-dark cycle and sample collection protocol for misaligned mice, subjected to a 6-hour phase advance 5 days prior to sacrifice. d) UMAP visualization of scVI latent embedding for all cells, colored by cell type. e) UMAP visualization of SMCs, colored by subtype. f) Number of cells per major cell type cluster. g) Number of cells per SMC subtype cluster.

In total, we generated 152,473 cells. After quality control for library size (minimum 1000 UMIs), mitochondrial UMI fraction (maximum 0.2), and doublet removal using Scrublet [25], 141,752 high-quality cells remained. To obtain a low-dimensional embedding, 1150 highly variable genes were detected (Methods) and used as input to scVI [26], with library size, Bmal1 knockout status, misalignment status, batch, sex, and time as covariates. Cells were clustered using the Leiden algorithm [27] with resolution 0.05, yielding 5 main clusters: 115,460 vascular smooth muscle cells (SMCs), 19,759 fibroblasts, 3,935 endothelial cells (ECs), 2,917 macrophages, and 631 T cells (Figures 1d, 1f). SMCs were further subclustered into 5 subtypes (Figures 1e, 1g). The majority of SMCs expressed canonical contractile markers, with variation primarily reflecting physical location: SMC0 (Hoxb5+, Hoxb6+) and SMC1 (Lgals3+, Hand2+) expressed markers of the aortic arch and ascending aorta [28, 29, 30], while SMC2 (Hoxa10+, Hoxc10+, Hoxa9+, Rgs5+) expressed markers of the descending thoracic aorta [28, 29, 30, 31]. Smaller subpopulations expressed markers of synthetic phenotypes: SMC3 expressed osteoblast [32, 33] (Runx2, Frzb) and chondrocyte [34] (Sox9) markers, while SMC4 expressed markers of multipotent SEM cells [35] (Ly6a, Ly6c1, Vcam1).

To estimate circadian expression parameters, approximate posterior distributions of gene temporal waveforms were inferred using Bayesian variational inference for each gene, cell type, and condition combination (Methods). Circadian cycling genes were identified by decomposing posterior waveforms using a Fast Fourier Transform (FFT) and computing the expected decrease in data likelihood when assuming the 24-hour component amplitude was zero (Methods). As a data quality check, we assessed whether the peak times of SMC cycling genes in our single-cell data aligned with peak times from bulk aorta transcriptomics, given that SMCs comprise most of the vessel wall. The posterior acrophases of SMC cycling genes aligned well with their peak times from bulk aorta generated by Zhang et al. [36] (Supplementary Figure 1a). As anticipated, core circadian clock genes demonstrated strong rhythms across all major cell types (Supplementary Figure 1c) and SMC subtypes (Supplementary Figure 1d). The acrophase distribution of cycling genes appeared bimodal, with peaks at dawn and dusk (Supplementary Figure 1b). Data from inducible Bmal1 deletion mice (Supplementary Figure 10) revealed very few cycling genes (Supplementary Figure 11), suggesting the vast majority of detected cycling in wild-type mice reflects regulation by the clock, either directly or indirectly. Altogether, these analyses provided confidence in both our data quality and our approach to calling cycling genes and estimating their parameters.

### Cell-type-specific daily cycling expression in the mouse aorta

To identify genes with robust daily cycling shared between sexes, we intersected male and female cycling gene lists within each cell type, requiring acrophases to be within 3 hours. This yielded 416 shared daily cyclers in SMCs, 117 in fibroblasts, 21 in ECs, and 5 in macrophages (Figure 2a). The vast majority of these cyclers appear to be regulated directly or indirectly by the molecular clock, as inducible Bmal1 deletion dramatically reduced the number of cycling genes in both SMCs and fibroblasts (Supplementary Figure 11a,b).

**Figure 2:**
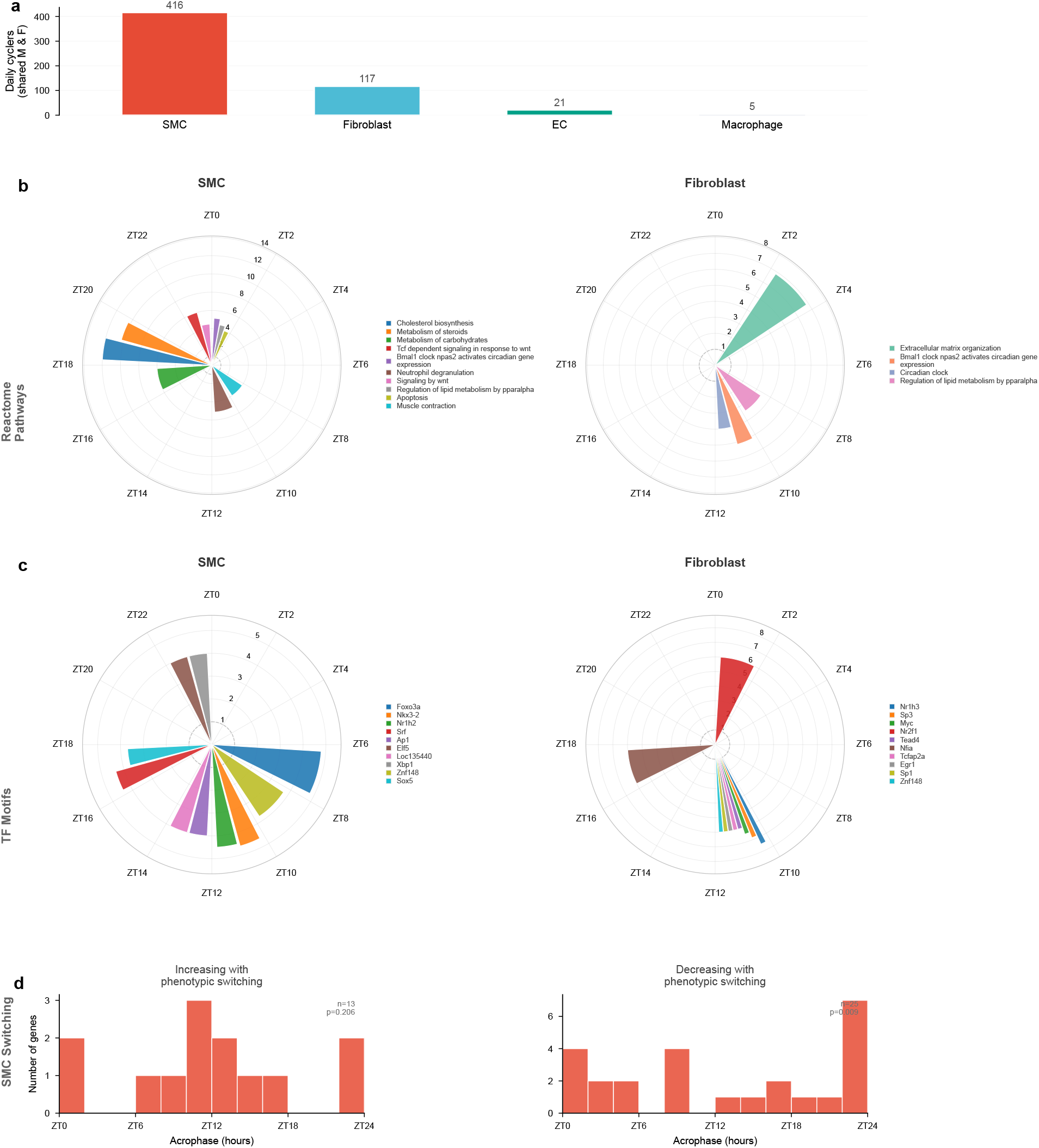
Cell-type-specific daily cycling expression in the mouse aorta. a) Number of daily cycling genes shared between males and females in each major cell type, requiring acrophases within 3 hours. b) Reactome pathway enrichment among daily cyclers in SMCs (left) and fibroblasts (right). Rose plots show enriched gene sets positioned by the mean acrophase of their member genes; radial extent indicates the number of member genes. c) Transcription factor motif enrichment among daily cyclers in SMCs (left) and fibroblasts (right), plotted as in (b). d) Acrophase distributions of SMC daily cyclers implicated in phenotypic switching. Left: genes that monotonically increase during the SMC-to-SEM transition. Right: genes that monotonically decrease. *n* and *p*-values from permutation enrichment tests for dusk (left) or dawn (right) peak times.

To identify pathways enriched among daily cyclers, we performed Reactome pathway enrichment analysis, restricting to gene sets whose members peak at similar times (Methods). In SMCs, the most strongly enriched pathways included cholesterol biosynthesis (peaking at dusk), metabolism of steroids, metabolism of carbohydrates, Wnt signaling, muscle contraction [41, 42, 43], PPAR*α*-regulated lipid metabolism, circadian clock, apoptosis, and degradation of the extracellular matrix (Figure 2b). As has been previously described [41], many features of aortic metabolism undergo circadian rhythms. In fibroblasts, daily cyclers were enriched for extracellular matrix organization (peaking at dawn), circadian clock, and PPAR*α*-regulated lipid metabolism (Figure 2b). EC and macrophage cycler counts were too low for robust pathway enrichment. We additionally performed transcription factor (TF) motif enrichment, identifying gene sets whose members share promoter motifs and peak at coordinated times. In SMCs, enriched motifs included FOXO3A, Nkx3-2, SRF, and AP1 (Figure 2c). In fibroblasts, enriched motifs included NR1H3, SP3, MYC, EGR1, SP1, and TEAD4, with most target genes peaking near ZT12 (Figure 2c). Male-specific and female-specific daily cycling analyses, including pathway enrichments, are shown in Supplementary Figures 2 and 3.

**Figure 3:**
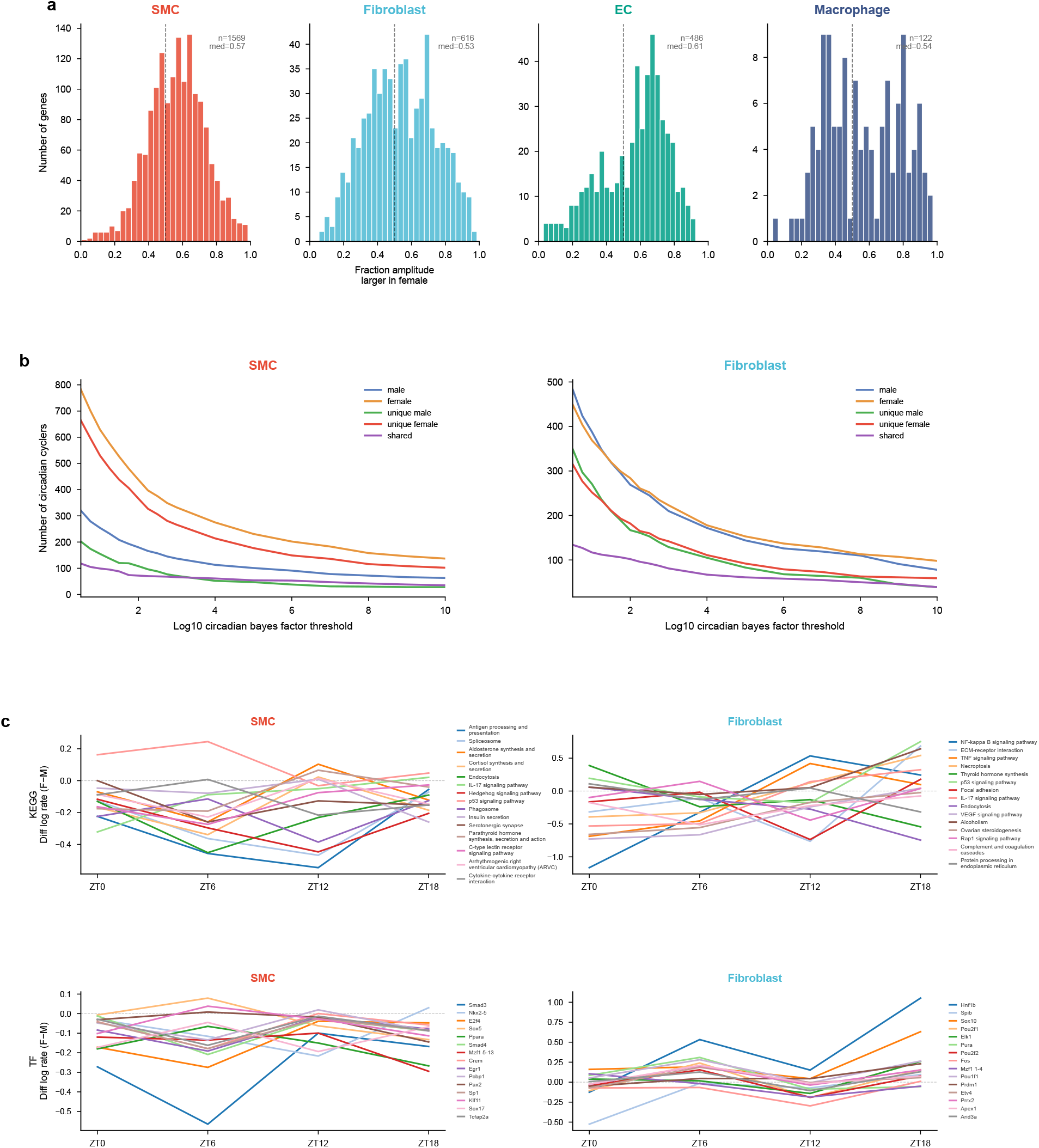
Enhanced rhythmicity of female aortic cell types. a) Distribution of the fraction of posterior amplitude samples larger in females than males, for genes cycling in either sex. Dashed line indicates 0.5 (no difference); median shown. Distributions shifted right indicate higher female amplitudes. b) Number of circadian cycling genes in males and females across log_10_ Bayes factor thresholds, after downsampling cells and pseudobulk UMI counts to match between sexes. SMCs (left) and fibroblasts (right). c) Sex-differential waveform enrichment analysis. Difference waveforms (female *−* male) for KEGG pathway gene sets (top) and transcription factor motif gene sets (bottom) in SMCs (left) and fibroblasts (right). Each line represents a gene set; values above zero indicate higher expression in females at that time point.

We next assessed whether daily cycling expression was relevant to SMC phenotypic switching in atherosclerosis. SMCs can dedifferentiate into stem cell and endothelial-like mesenchymal (SEM) cells, which are multipotent and can differentiate into macrophage-like and fibroblastlike cells [35, 37, 38, 39]. Daily cyclers (shared between males and females) in SMCs that monotonically increase from SMC to SEM tended to peak at dusk (ZT12), while those that decrease from SMC to SEM tended to peak at dawn (ZT0) (Figure 2d, Supplementary Figures 2 and 3 for male and female specific analyses). Though circadian gating of cell fate decisions has been observed in numerous contexts [69, 70, 71], it remains unknown whether the clock gates SMC phenotypic switching.

### Enhanced rhythmicity of female aortic cell types

Among shared cycling genes between male and female cell types, female circadian gene expression was distinguished by higher amplitude rhythms (Figure 3a). Moreover, among cycling genes common to both sexes, acrophases tended to peak modestly earlier in females, consistent with previous observations [21]. To determine whether female cell types have more cyclers, we controlled for cell count and library size discrepancies by downsampling male and female cells and pseudobulk UMI counts to match within each cell type before rerunning our Bayesian regression and hypothesis testing pipeline (Methods). After this adjustment, female SMCs retained nearly twice as many cycling genes as males across Bayes factor thresholds, while fibroblasts showed comparable numbers between sexes (Figure 3b). These results suggest sex-dependent rhythmicity is cell-type-specific. Macrophages additionally demonstrated stronger female rhythmicity, though this result should be interpreted cautiously given the low cell count. Notably, females retained more cycling genes than males even after inducible Bmal1 deletion (Supplementary Figure 11a,b). Moreover, among wild-type cyclers in SMCs, expression became more similar between sexes after knockout, suggesting that a substantial portion of sex-specific cycling expression is mediated through the circadian clock (Supplementary Figure 11c).

To identify pathways and transcription factor networks with sex-differential temporal expression, we performed a two-group waveform enrichment analysis, computing the difference waveform (female *−* male) for each gene set and testing whether it deviates from zero at each time point (Methods). In SMCs, gene sets involved in antigen processing and presentation, spliceosome, protein processing in the endoplasmic reticulum, endocytosis, and estrogen signaling showed lower expression in females, particularly at ZT6 and ZT12 (Figure 3c). Notably, the lower female expression of the KEGG estrogen signaling pathway reflects its gene composition: the pathway members detected in our data are predominantly heat shock protein chaperones (Hspa1a, Hspa1b, Hsp90aa1, Hspa8, Hsp90b1), which are known to have higher expression in male SMCs rather than differences in estrogen receptor signaling itself. Among transcription factor networks, targets of SMAD3, Nkx2-5, and SMAD4 showed sex-differential temporal patterns, with lower female expression concentrated at ZT6 (Figure 3c). In fibroblasts, NF-*κ*B signaling, ECM-receptor interaction, and TNF signaling gene sets showed the largest amplitude sex differences (Figure 3c). Extended Reactome and GO Biological Process waveform enrichment results are shown in Supplementary Figure 4.

### Circadian misalignment disrupts proteostasis and cell-type entrainment in the aorta

We first examined the effects of acute misalignment in male mice (female counterparts are presented in Supplementary Figures 7–9). After a 6-hour phase advance, the number of cycling genes decreased across all cell types, even after controlling for cell count and library size differences (Supplementary Figure 5). Among cycling genes common to both conditions, amplitudes were generally decreased (Figure 4a) and acrophases were generally delayed in misaligned mice, indicating incomplete entrainment to the new LD cycle (Figure 4b).

**Figure 4:**
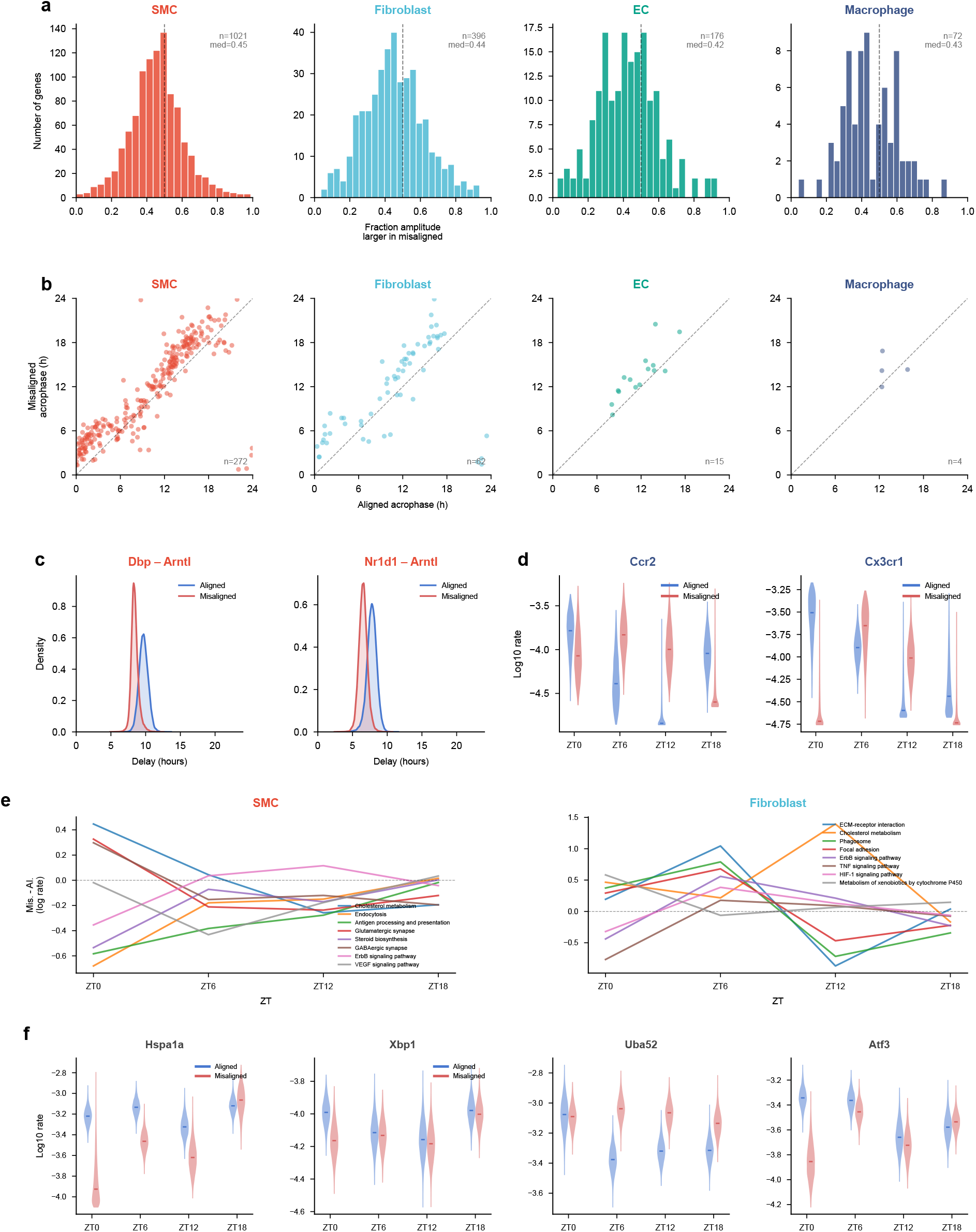
Circadian misalignment disrupts proteostasis and cell-type entrainment in the aorta. a) Distribution of the fraction of posterior amplitude samples larger in misaligned than aligned mice, for genes cycling in either condition. Distributions shifted left of 0.5 indicate reduced amplitudes after misalignment. b) Acrophase scatter plots comparing aligned (x-axis) and misaligned (y-axis) peak times for cycling genes shared between conditions. Dashed line indicates identity (perfect entrainment); points above the line indicate delayed acrophases. *n*, number of shared cycling genes. c) Posterior distributions of the phase delay between Dbp and Arntl (left) and Nr1d1 and Arntl (right) in SMCs, comparing aligned and misaligned conditions. Compression of the delay indicates altered intracellular clock architecture. d) Posterior waveforms of macrophage Ccr2 (left) and Cx3cr1 (right) in aligned and misaligned conditions. Violin plots show posterior log_10_ transcript proportion estimates at each time point; horizontal lines indicate posterior means. e) KEGG pathway waveform enrichment showing the difference in log_10_ expression rate (misaligned *−* aligned) at each time point for SMCs (left) and fibroblasts (right). Values below zero indicate decreased expression after misalignment. f) Posterior waveforms of representative proteostasis-related genes in SMCs, comparing aligned and misaligned conditions. Violin plots as in (d).

The central clock in the suprachiasmatic nucleus (SCN) is reported to re-entrain to a 6-hour phase advance over approximately 5 days in C57BL/6J mice [144], meaning the master clock had likely adapted to the new LD cycle by the time of tissue collection. The incomplete reentrainment of vascular cell types at this time point is consistent with internal desynchrony between the peripheral vasculature and the SCN-driven central timing system. Similar SCN– peripheral clock dissociation has been observed in other peripheral clocks [145] following LD-cycle shifts, where peripheral clocks re-entrain more slowly than the SCN [145, 146]. This transient internal misalignment may extend across peripheral organs and is implicated in the adverse physiological consequences of jet lag and shift work.

Beyond desynchrony between the central and peripheral clocks at the tissue level, misalignment also appeared to alter the intracellular timing relationships among clock genes within SMCs. Restricting to clock genes whose posterior acrophase 95% interval was within 2 hours in both conditions, we found that Dbp and Nr1d1 — both direct transcriptional targets of BMAL1:CLOCK — compressed their phase delay relative to Arntl by approximately 1.3 and 1.2 hours, respectively (Figure 4c). This suggests that even within individual cell types, the internal architecture of the clock may be rearranged during acute misalignment. Acute misalignment also provoked gene expression patterns consistent with altered timing of chemoattraction of myeloid cells. While most shared cycling genes shifted acrophases slightly toward the new LD cycle, macrophage Ccr2 and Cx3cr1 — which mediate rhythmic myeloid chemoattraction [4, 89, 90, 91] — showed acrophase delays of approximately 6 hours, suggesting minimal adaptation to the new LD cycle (Figure 4d). However, this should be interpreted cautiously given the low cell counts and shallow sequencing depth in macrophages. Moreover, whether these changes extend to monocytes remains unknown.

The most striking effect of misalignment in male SMCs was disruption of proteostasis gene expression (Figure 4e,f). Hspa1a, a major chaperone involved in heat shock response and the unfolded protein response, was substantially downregulated, particularly at ZT0 (Figure 4f). Additional chaperones — including Hspa1b, Dnajb1, Dnajb4, Dnaja1, Hsph1, and Ankrd1 — were similarly downregulated (Supplementary Figure 6c), as was Paip2b, an inhibitor of mRNA translation [107]. Consistent with disrupted proteostasis, Xbp1, a key regulator of the unfolded protein response implicated in preventing neointimal hyperplasia [105], was downregulated at ZT0 (Figure 4f). Conversely, Uba52, which targets proteins for degradation [108, 109], was strongly upregulated (Figure 4f), and gene sets involved in transcription and ribosome biogenesis showed decreased mesors (Figure 4e; Supplementary Figure 6a,b) — possibly reflecting compensatory efforts to reduce potential unfolded protein burden. Atf3, which has been described to inhibit SMC phenotypic switching, was substantially decreased at ZT0 with an acrophase delay of approximately 7 hours (Figure 4f), suggesting acute misalignment may additionally predispose SMCs to phenotypic switching. Female-specific responses to acute misalignment are presented in Supplementary Figures 7–9, which show broadly similar patterns of proteostasis disruption and incomplete entrainment to the new LD cycle.

### Sex-specific and shared vascular and thrombotic responses to acute circadian misalignment

To determine whether the observed transcriptomic changes of acute misalignment were physiologically relevant, a series of assays were conducted to assess vascular function in C57BL/6J mice. Three days following a 6-hour phase advance, male misaligned mice at ZT12 exhibited significantly decreased urinary nitrate concentration normalized to urinary creatinine (p < 0.05; Figure 5a). Female mice also exhibited a decrease in urinary nitrate concentration at ZT12, although this did not reach statistical significance (p = 0.0649). Furthermore, both male and female mice exhibited significantly increased vascular permeability at ZT12, as determined by Evans blue extravasation into aortic tissue five days post-misalignment (Figure 5b). Collectively, these findings may indicate impaired endothelial function following acute misalignment.

**Figure 5:**
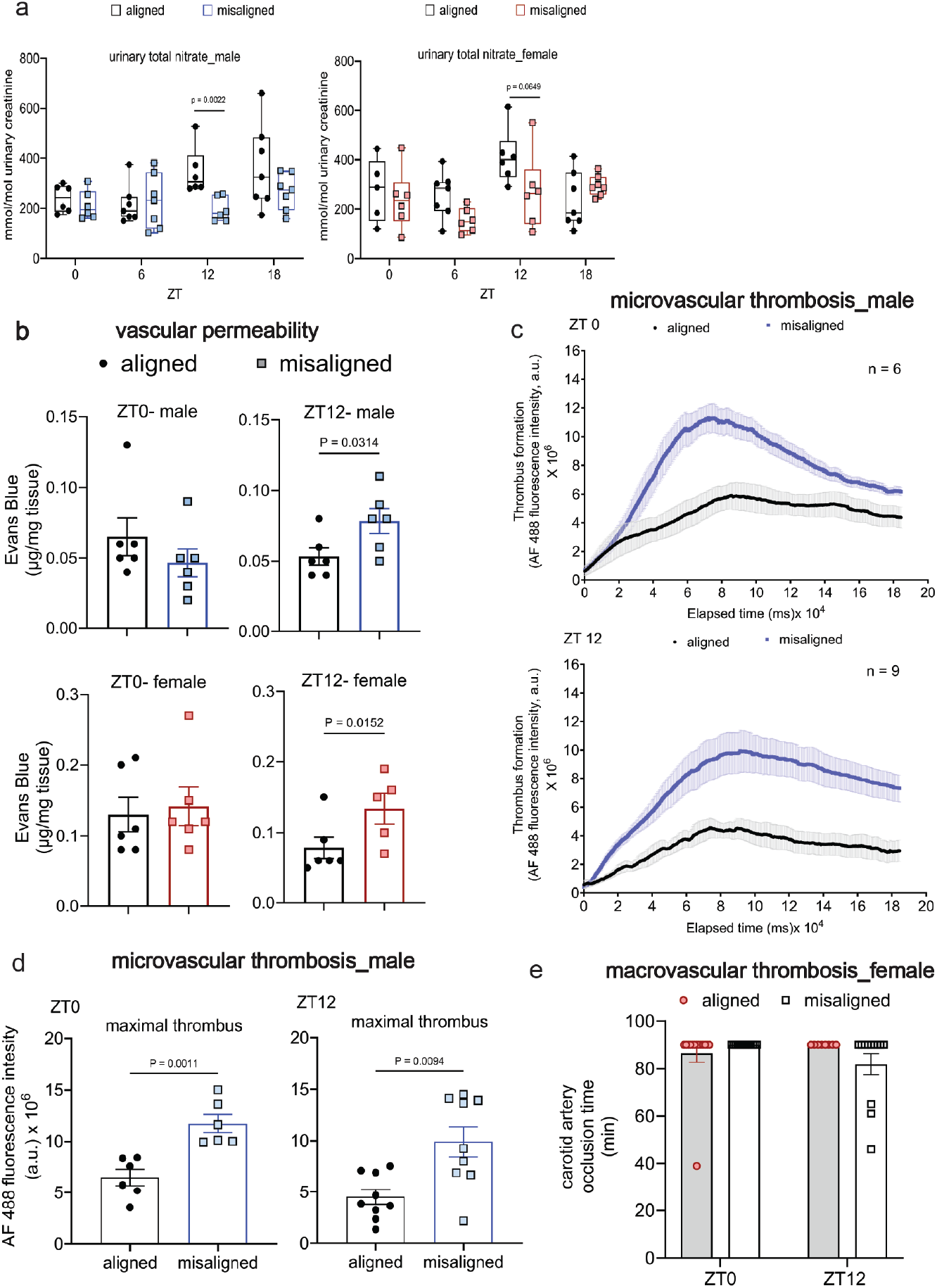
Acute circadian misalignment alters vascular permeability and thrombus formation in a sex- and time-dependent manner. a) Urinary total nitrate concentrations, normalized to urinary creatinine, in aligned and misaligned male (left) and female (right) mice measured at ZT0, ZT6, ZT12, and ZT18. b) Vascular permeability assessed by Evans Blue extravasation in male (top) and female (bottom) mice at ZT0 and ZT12 following acute circadian misalignment. c) Representative time-course measurements of microvascular thrombus formation in aligned and misaligned male mice at ZT0 (top) and ZT12 (bottom), quantified by fluorescence intensity following photochemical injury. d) Maximum microvascular thrombus fluorescence intensity in aligned and misaligned male mice at ZT0 (left) and ZT12 (right). e) Time to carotid artery occlusion following vascular injury in aligned and misaligned female mice at ZT0 and ZT12. Data are presented as mean *±* SEM, with individual data points shown.

Given the observed alterations in endothelial function and the concomitant dysregulation of endothelial cell genes implicated in thrombosis, we next investigated whether acute misalignment was associated with a functional prothrombotic phenotype in both male and female mice. Acute misalignment significantly increased microvascular thrombus formation in male mice at both ZT0 and ZT12 (p < 0.05) five days post-misalignment, as assessed using laser-induced thrombosis and intravital microscopy of the cremaster muscle (Figures 5c and 5d). In contrast, no significant increase in thrombosis was observed in female mice at either ZT0 or ZT12 following photochemical vascular injury of the carotid artery (Figure 5e).

## Discussion

Circadian clock disruption has a notable influence on cardiovascular health, yet the cell-type-specific molecular basis by which misalignment with the light-dark cycle affects the vasculature remains poorly understood. Bulk RNA-sequencing circadian time courses of the aorta have defined key vascular functions that cycle with circadian rhythms [41], but cannot resolve cell-type-specific contributions. Here, we used scRNA-seq to generate a cell-type-specific circadian atlas of the mouse aorta, characterize sex differences in rhythmicity, and define the transcriptomic consequences of acute circadian misalignment. We then assessed vascular function in vivo to determine whether a 6-h phase advance produced functional impairment 5 days after the shift.

Our atlas reveals that the extent of circadian transcription varies markedly across vascular cell types. SMCs and fibroblasts harbor hundreds of cycling genes, while ECs and macrophages show far fewer — likely reflecting both biological differences in clock output and lower statistical power from smaller cell populations. Pathway enrichment among SMC cyclers highlighted coordinated rhythms in cholesterol biosynthesis, muscle contraction, extracellular matrix organization, and lipid metabolism, consistent with prior bulk aorta studies [41, 42, 43]. An intriguing finding was the temporal coordination of genes implicated in SMC phenotypic switching: genes that increase during the SMC-to-SEM transition peaked at dusk, while those that decrease peaked at dawn. Although circadian gating of cell fate decisions has been observed in other contexts [69, 70, 71], whether the clock gates SMC phenotypic switching remains an open question warranting functional investigation.

Comparing males and females, we found that sex differences in rhythmicity were cell-type-specific rather than uniform. Female SMCs exhibited higher amplitude rhythms and nearly twice as many cycling genes after controlling for cell counts and library sizes, whereas fibroblasts showed comparable rhythmicity between sexes. These findings are consistent with previous observations that females exhibit more robust circadian characteristics [21]. However, as the study was not specifically powered to compare circadian effects across sex and vascular cell type, the absence of a detectable female-associated effect in other vascular cell types should not be interpreted as evidence that such effects are absent. Rather, the findings suggest that sex-associated circadian responses may vary across vascular cell types, a possibility that should be examined in appropriately powered studies. Future work integrating sex-hormone perturbations with cell-type-resolved circadian profiling may help distinguish the contributions of hormonal signalling, cell-intrinsic clock properties, and differences in effect size or detectability across cell types.

The most notable findings of our study concern the effects of acute circadian misalignment, which appeared to disrupt circadian organization at multiple levels. At the tissue level, vascular cell types showed incomplete re-entrainment to the new LD cycle 5 days after the phase advance. Given that the central clock in the SCN is known to re-entrain within this timeframe [144], this implies internal desynchrony between central and peripheral clocks — a state proposed to underlie many adverse health consequences of jet lag and shift work [6]. At the intracellular level, the phase relationships among clock genes were altered: Dbp and Nr1d1 altered their phase delay relative to Arntl in males and females, suggesting the internal architecture of the molecular clock may be rearranged during acute misalignment. At the intercellular level, tissue-resident macrophage expression of Ccr2 and Cx3cr1 — which participate in signaling axes that mediate rhythmic myeloid recruitment to the vasculature [4, 89, 90, 91] — showed minimal adaptation to the new LD cycle, in contrast to the partial re-entrainment observed in endogenous vascular cell types. This differential adaptation may disrupt the coordinated timing between macrophage and vascular cell type signaling, though the functional consequences of such intercellular desynchrony remain to be tested.

Beyond these timing disruptions, the most striking molecular consequence of misalignment in SMCs, the predominant cell type present in aortic tissue, was disruption of proteostasis. Multiple chaperone and stress-response genes were substantially downregulated, including key components of the HSP70 system (Hspa1a, Hspa1b, Dnajb1, Dnajb4, Dnaja1, Hsph1) and regulators of the unfolded protein response (Xbp1, Ankrd1). Downregulation of the translational repressor Paip2b may exacerbate this burden by permitting continued protein synthesis. Conversely, upregulation of Uba52, which targets proteins for proteasomal degradation [108, 109], and decreased mesors of gene sets involved in transcription and ribosome biogenesis may reflect compensatory efforts to reduce the accumulation of misfolded proteins. Disrupted proteostasis has been implicated in vascular pathology, including ER stress-driven neointimal hyperplasia [105] and atherosclerotic plaque instability. Whether acute misalignment indeed increases the unfolded protein burden, and whether these compensatory responses are sufficient to maintain proteostasis, will require functional follow-up. Additionally, the substantial downregulation of Atf3, which has been described to inhibit SMC phenotypic switching, suggests that acute misalignment may predispose SMCs to dedifferentiation — a possibility reinforced by the coordinated temporal patterns of switching genes observed in aligned mice.

Importantly, the molecular disturbances observed were accompanied by functional vascular abnormalities several days after the phase advance compared to controls. Both sexes exhibited compromised vascular barrier integrity, as evidenced by increased aortic Evans blue extravasation at ZT12 five days post misalignment. Urinary nitrate was reduced significantly in males three days post misalignment and showed a similar, though non-significant, decrease in females, consistent with impaired nitric oxide bioavailability and potential endothelial dysfunction. These observations are biologically plausible given the role of endothelial nitric oxide signaling in maintaining vascular permeability, vascular homeostasis, and resistance to thrombosis [136, 137].

However, the thrombotic consequences of misalignment were sex-specific: males displayed enhanced microvascular thrombus formation at both ZT0 and ZT12, whereas females showed no significant increase in carotid thrombosis. This divergence suggests that endothelial impairment alone may be insufficient to determine thrombotic susceptibility. Sex-dependent differences in haemostatic pathways, vascular injury responses, nitric oxide signalling, or sex hormone status may modify the downstream response to circadian disruption. It is also possible that female circadian resilience extends beyond the central clock to vascular and haemostatic systems, thereby attenuating disruption-induced changes in nitric oxide signalling and thrombotic susceptibility. However, the present data cannot distinguish these possibilities. Notably, the use of distinct thrombosis models in males and females precludes a direct quantitative comparison of thrombotic susceptibility between sexes. Nevertheless, these findings identify acute circadian misalignment as sufficient to induce shared vascular dysfunction while revealing a potentially male-biased prothrombotic phenotype, reinforcing the importance of considering sex as a biological variable in studies of circadian cardiovascular risk [138, 139].

Our study has several limitations. While we collected many SMCs, allowing high-confidence circadian parameter estimation, we captured relatively few ECs and macrophages. The number of cyclers in these populations is likely an underestimate, and macrophage-specific findings — including the intercellular timing observations — should be interpreted cautiously given the low cell counts and shallow sequencing depth. Additionally, our scRNA-seq data capture transcript-level changes only; the proteostasis disruption we infer from gene expression patterns requires biochemical validation. Our study also captures a single time point after misalignment (5 days post-advance), and the trajectory of adaptation — whether these effects resolve, persist, or worsen — remains unknown.

The in vivo findings should also be interpreted within several methodological constraints. Urinary nitrate normalised to creatinine provides an indirect systemic measure of nitric oxide metabolism and does not establish reduced nitric oxide production specifically within aortic ECs, while Evans blue extravasation indicates increased albumin-associated dye accumulation without defining the underlying mechanism of vascular leak. In addition, the use of distinct thrombosis models and vascular beds in males and females prevents a direct quantitative comparison of sex-specific thrombotic susceptibility. Finally, functional outcomes were assessed at limited circadian time points following a single acute phase advance; future studies should establish whether these changes represent sustained vascular dysfunction, altered rhythmicity, or a transient response to misalignment.

Altogether, this atlas provides a resource for understanding how the circadian clock organizes vascular gene expression across cell types and sexes, and how acute misalignment disrupts this organization at multiple scales — from intracellular clock architecture to intercellular coordination to proteostatic homeostasis. These molecular changes were accompanied by increased vascular permeability in both sexes, reduced urinary nitrate in males, and enhanced thrombus formation in male mice, demonstrating that a transient phase advance can produce persistent vascular dysfunction during the early re-entrainment period, consistent with the overall dimorphism in cardiovascular risk to which the clock may contribute. These findings may help explain why circadian misalignment elevates cardiovascular risk and identify proteostasis, alongside endothelial dysfunction and sex-dependent thrombotic responses, as candidate processes linking circadian disruption to vascular disease. Future work should define the mechanisms connecting these cell-type-specific transcriptional changes to vascular function and establish their persistence under repeated or chronic circadian misalignment.

## Methods

### Inducible deletion of the Bmal1 gene in Bmal1fl/fl:CAGGcreERt2/+ mice

At 8 weeks of age, Bmal1 was inducibly deleted in Bmal1fl/fl:CAGGcreERt2/+ mice by administering tamoxifen via oral gavage (100 mg/kg of body weight) for 5 days. Protein expression loss of Bmal1 was confirmed via Western blot (Supplementary Figure 10).

### Aligned and acute misalignment conditions in C57BL/6J mice

For the aligned conditions, 12-week-old C57BL/6J and Bmal1fl/fl:CAGGcreERt2/+ male and female mice from Jackson Laboratory were entrained to a 12:12 LD cycle (12 hours of light, followed by 12 hours of darkness) for 2 weeks in circadian boxes with ventilation (20–22*◦*C, relative humidity *∼*50%) and fed ad libitum. At ZT0, ZT6, ZT12 (lights off), or ZT18 mice were sacrificed (2 per time point) via CO_2_ exposure. For the misaligned conditions, 12-week-old C57BL/6J male and female mice from the Jackson Laboratory were entrained to a 12:12 LD cycle (12 hours of light, followed by 12 hours of darkness) for 2 weeks in circadian boxes with ventilation (20–22*◦*C, relative humidity *∼*50%) and fed ad libitum. Mice were then phase advanced 6 hours by changing the lighting schedule. Then, 5 days after the phase advance, mice were sacrificed at ZT0, ZT6, ZT12 (lights off), or ZT18 (2 per time point).

### Urinary nitrate colorimetry and mass spectrometric analysis of urinary creatinine

On the third day following the 6-hour phase advance, mice were briefly transferred to metabolic cages at ZT0, ZT6, ZT12, or ZT18. Urine was collected into Eppendorf tubes and stored at *−*20*^◦^*C until analysis. Urinary nitrate concentrations were measured using a nitrate/nitrite colorimetric assay kit (Cayman Chemical, cat. no. 780001) according to the manufacturer’s instructions. Urinary nitrate concentrations were normalized to urinary creatinine concentrations. Urinary creatinine was quantified by UPLC-MS/MS using positive-mode electrospray ionization (ESI) and multiple reaction monitoring (MRM) according to the following method [140]. For sample preparation, 1 mL of stable isotope-labeled internal standard ([d3]-creatinine, 2.5 *µ*g/mL in 3% H_2_O/acetonitrile) was added to 5 *µ*L of mouse urine, and the resulting solution was diluted 10-fold. Samples were transferred to autosampler vials, and 5 *µ*L was injected onto the UPLC-MS/MS system. Chromatographic separation was performed using a Waters ACQUITY UPLC system equipped with a Waters XBridge BEH HILIC column (2.1 *×* 50 mm, 2.5 *µ*m particle size). Mobile phase A consisted of 100% acetonitrile, while mobile phase B consisted of 5 mM ammonium formate (pH 5.7). The mobile phases were delivered at a flow rate of 350 *µ*L/min under isocratic conditions with 12% mobile phase B. Detection was performed using a Waters Xevo TQS triple quadrupole mass spectrometer operated in positive-mode ESI and MRM mode. The ion transitions monitored were m/z 114 *→* 86 for creatinine and m/z 117 *→* 89 for the [d3]-creatinine internal standard. Quantitation was performed by calculating the peak area ratio of creatinine to the internal standard using TargetLynx software (version 4.1), with results normalized to sample volume.

### Fluorescence intravital microscopy of laser-induced thrombus formation in male mice

Experimental procedures were conducted as previously described by Yu and colleagues [142]. All experiments were conducted at either ZT0 or ZT12. In brief, approximately 15-16-week-old male mice post control or misaligned conditions, were anesthetized via intraperitoneal injection of ketamine (125 mg/kg) and xylazine (12.5 mg/kg) and maintained at 37*◦*C using a thermal pad. The scrotum was incised, the cremaster muscle exteriorized, and pinned onto the microscopy stage. The tissue was continuously superfused with bicarbonate-buffered saline at 37*◦*C, aerated with 95% N_2_ and 5% CO_2_. Circulating platelets were fluorescently labelled by intravenous administration of rat anti-mouse CD41 (20 *µ*g/mL; BD Pharmingen, #553847) and Alexa Fluor 488 chicken anti-rat IgG (180 *µ*g/mL; Life Technologies, #A21470), diluted in PBS. The antibody mixture (4 *µ*L/g body weight) was administered via the jugular vein through the pectoral muscle. Arteriolar injury within the cremaster muscle was induced using a fluorescence resonance energy transfer/fluorescence recovery after photobleaching (FRET/FRAP) photoablation system. A 440 nm laser, focused through the microscope objective, parfocal with the focal plane, was directed at the vessel wall. Injury was induced using a single pulse from a nitrogen dye laser at *∼*70% power (mW). Sequential thrombi were generated either upstream of the prior injury site or in different arterioles within the same cremaster preparation. Image acquisition was performed on an Olympus BX61WI microscope controlled by SlideBook 6.0 imaging software (Intelligent Imaging Innovations). Wide-field fluorescence microscopy employed a 2-Galvo high-speed wavelength changer with a 300-W xenon light source and excitation filters (360, 480, 575, and 655nm). Signal intensity was enhanced up to 1000-fold using a Hamamatsu image intensifier. High-resolution images (1390 × 1024 pixels) were captured with a Hamamatsu C9300-201 CCD camera, enabling both bright-field and fluorescence imaging. Image analysis was performed using SlideBook 6.2 software, incorporating modules for image reconstruction, deconvolution, statistical analysis, and volumetric quantification. Mean background fluorescence was determined for each image as the average pixel intensity within a defined region of the vessel upstream of the injury site, where circulating fluorescent antibody was the primary contributor. Thrombus area was defined as the total number of pixels with fluorescence intensity exceeding the maximal upstream background fluorescence measured during image acquisition. Background correction was applied by multiplying the mean upstream fluorescence intensity by the thrombus area at each time point. Thrombus volume was calculated as the total number of voxels across all image planes with fluorescence intensity above the maximal background level.

### Photochemical vascular injury induced thrombosis in female mice

As previously described [142], control or misaligned mice were anesthetized by intraperitoneal injection of ketamine (125 mg/kg) and xylazine (12.5 mg/kg) and maintained at 37*◦*C using a thermal pad. Following a midline cervical incision, the right common carotid artery was identified and a Doppler flow probe was applied (Model 0.5 VB, Transonic Systems, Ithaca, NY, USA). The probe was connected to a flowmeter and data acquisition was achieved using the Powerlab computer program (AD Instruments, CO, USA). Rose Bengal (10 mg/mL in saline; 50 mg/kg body weight) was administered via the jugular vein, and thrombosis was induced using a 1.5 mW green laser (540 nm) at the desired site of injury on the exposed carotid artery from a distance of 5 cm. Blood flow was monitored for up to 90 minutes, and occlusion time was defined as the point at which blood flow reached zero for at least 1 minute. All experiments were carried out at either ZT0 or ZT12.

### Evans blue assay for vascular leakiness

Control and circadian-misaligned mice were assessed for vascular permeability (plasma extravasation) on day 6 following misalignment at ZT0, ZT6, ZT12, and ZT18. Male and female mice were anesthetized with 5% isoflurane in an induction chamber. Once a surgical plane of anesthesia was achieved, anesthesia was maintained via nose cone at 3% isoflurane. Evans blue dye was administered by retro-orbital injection (20 mg/kg in sterile saline), after which mice were returned to their home cages. After 20 mins, the mice were euthanized by cervical dislocation. The aortae of mice were quickly isolated and the perivascular fat tissue removed. To measure the Evans blue in tissue [143], each of the aortae were placed in an Eppendorf tube with 200 µL formamide for 48 h at room temperature to extract the Evans blue. Then 50 µL of the Evans blue-infused formamide for each tissue was added to a 96-well plate, using pure formamide as blanks. Absorbance was measured at 620 nm (OD620) using a microplate reader, corresponding to the absorbance maximum of Evans blue. The tissue remaining post infusion was dried at 55*◦*C overnight to enable weight/dry weight quantification of the extent of vascular leakiness. Data are presented as Evans blue dye in µg/mg tissue.

### Aorta collection and scRNA-seq data generation

For each mouse in each condition, whole aorta was dissected and cut into small pieces (*∼*1 mm), and cells were disassociated with 1.5 mL of enzyme cocktail (DNase–120 U/mL (Worthington, #LS006331), Liberase TM–4 U/mL (Roche, #05401127001), and hyaluronidase–60 U/mL (Sigma-Aldrich, #H3506)) were added to a petri dish at 37*◦*C for 40 mins. A 1 mL pipette was used to aspirate the tissue every 10 minutes during incubation to aid dissociation of aortic cells. Cell supernatant was filtered through a 40 *µ*m strainer and washed with RPMI1640 (Gibco, #1187-085) containing 10% fetal bovine serum (FBS, HyClone, #SH30071.03) to inactivate the enzyme cocktail. Residual red blood cells were lysed by incubating the cell suspension with Red Blood Cell Lysing Buffer Hybri-Max (Sigma-Aldrich, #R7767) at RT for 1 min. The cells were washed two more times to remove debris with FACS buffer (FBS 2%), EDTA (5 mM, Invitrogen, #15575-038), HEPES (20 mM, Gibco, #15630-080), sodium pyruvate (1 mM, Gibco, #11360-070) in 1x PBS (Gibco, #14190-136), and resuspended in DMEM/F12 (Gibco, #11320-033) media containing 10% FBS for further analysis. Cells from mice sacrificed at the same time point were pooled to form single-cell suspensions. Single cells were then captured and barcoded using the 10X Genomics Chromium platform and sequenced using an Illumina NovaSeq S2 flow cell. Cell barcode detection, read alignment, and transcript quantification were performed using the 10X Genomics Cell Ranger pipeline.

### Cell type marker gene detection procedure

Cell type markers used as input for low-dimensional embedding estimation were selected via a three-step procedure yielding 1969 marker genes.

First, markers whose variation defines major cell types were selected via a prior knowledge-based procedure. This was performed because preliminary clustering suggested high cell type imbalance of our dataset, with a disproportionate amount of SMCs and fibroblasts. As such, markers of rarer cell types (e.g. T cells) may not be identified using traditional methods based on detecting highly variable genes. Based on our prior work described in Pan et al. [35], we anticipate SMCs, fibroblasts, ECs, macrophages, and T cells. Based on this work, we selected several highly expressed and highly specific markers for each major cell type. We selected Myh11 and Acta2 for SMCs; Serpinf1 and Serping1 for fibroblasts; Vcam1 and Pecam1 for endothelial cells; Cd68, Cd14, Cd16, and Cd64 for macrophages; and Cd4, Cd8, Cd3g, and Cd3d for T cells. For each major cell type, we then identified cells belonging to that cell type based on the transformed expression of their corresponding prior knowledge marker genes. Expression was transformed as follows:

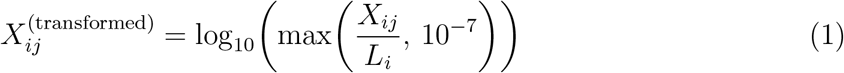

Where *X_ij_* is the UMI count of gene *j* in cell *i* and *L_i_* is the library size of cell *i*. Cells with transformed expression greater than *−*3 for any cell type marker were assigned to the corresponding cell type. This led to 119,817, 18,547, 3,632, 525, 185 cells roughly labeled as SMCs, fibroblasts, ECs, macrophages, and T cells. 5498 cells remained unlabeled.

Second, we identified markers with highly correlated expression to the prior knowledge cell type markers after the rough cell assignments. To identify these markers, the log_10_ pseudobulk relative expression of all genes was computed within each rough cell type. Cell type markers were then identified as those with log_10_ pseudobulk relative expression of at least *−*4 (i.e. 1 in 10,000 transcripts in pseudobulk) and with log_10_ pseudobulk relative expression larger than all other cell types by at least 0.5. This yielded 114, 113, 101, 228, 93 additional markers for SMCs, fibroblasts, ECs, macrophages, and T cells, respectively.

Third, we identified markers with high variability within cell types. To identify these marker genes, we first limited ourselves to genes that were sufficiently highly expressed. We decided cell type marker genes should be expressed at relative proportions of 1 in 10,000 transcripts in the cell types they mark. Moreover, there should be at least 30 cells observed for each cell type. Given this and given the median library size of 12,553 UMIs in our dataset, cell type marker genes should have at least 37 pseudobulk UMI for the cell type of interest (i.e. 10*^−^*^4^ *×* 30 *×* 12553). Using this set of genes, we next restricted genes to those that had highly variable expression. Using the transformed expression counts, we computed the mean and variance of each genes’ transformed expression. After fitting a nonparametric kernel (Python statsmodels implementation of Nadaraya-Watson, using bandwidth of 0.1 and default values for all other parameters) to describe the mean-variance relationship, genes with large variances were detected as those with Pearson residuals larger than 0.5.

### Bayesian temporal regression model

#### Likelihood model

A Bayesian ANOVA regression model was fit to describe gene waveforms and detect genes with circadian rhythms. Our model was fit independently for each cluster and condition combination. As input, our model requires an *n × p* transcript count matrix **X**, where *n* is the number of cells and *p* is the number of genes. For gene *j* in cell *i* with library size *L_i_* and cell phase Θ*_i_* (where Θ*_i_ ∈ {*1*,…, q}* discrete time points), the unique molecular identifier (UMI) count *X_ij_* was assumed to follow a Negative Binomial (NB) distribution:

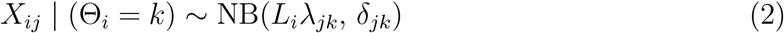

Where *k* is the discrete index of the time the cell was collected at,

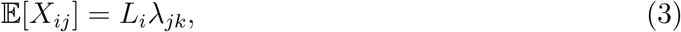

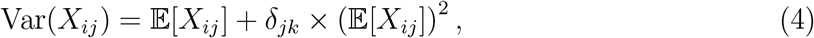

and

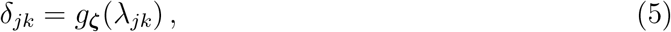

and where *g**_ζ_***(*λ_jk_*) is a deterministic polynomial function parameterized by ***ζ*** (shared by all cells and genes) describing the relationship between transcript proportion *λ_jk_* and the dispersion *δ_jk_*. Details on the estimation of ***ζ*** can be found in the “Estimating the transcript proportion – dispersion relationship” section below.

### Prior knowledge of gene parameters

Prior knowledge about *λ_jk_* is specified as a transformed Beta distribution:

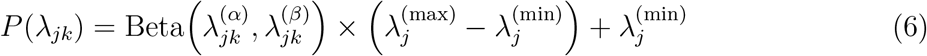

where 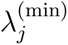 and 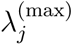 denote the minimum and maximum possible values for 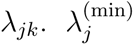 and 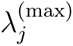 are set as follows. For each gene we compute the pseudobulk relative proportion (i.e. 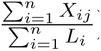). Assuming relatively balanced numbers of cells from each collection time point, the pseudobulk relative proportion is a reasonable estimate of each genes’ mesor. Moreover, based on analysis of past scRNA-seq circadian datasets, the largest observed fold changes of cycling genes from mesor to peak are around 10-fold (i.e. 1 in log_10_ space), while most have fold changes of around 2-fold. As such, we define 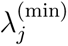 and 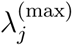 to account for fold changes of around 30-fold to be conservative (1.5 in log_10_ space):

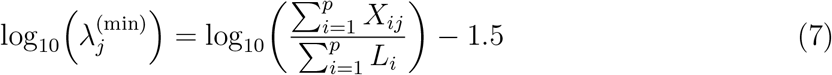

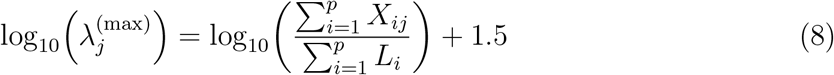

By default, Tempo sets 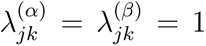, which assumes a non-informative prior over the domain of possible *λ_jk_* values.

### Approximate posterior inference

Using our prior knowledge of the gene parameters and the observed data, we seek the following joint posterior distribution of our gene parameters:

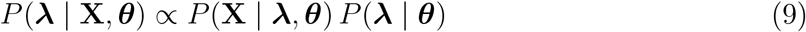

where ***λ*** is a *p × q* dimensional matrix containing the transcript proportion parameters for each gene at each time point, and

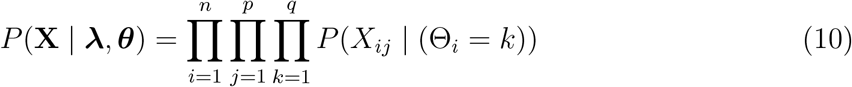

and

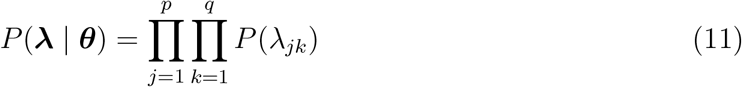

No known analytic solution exists to *P* (***λ*** *|* **X***, **θ***) *∝ P* (**X** *| **λ**, **θ***) *P* (***λ*** *| **θ***). Moreover, asymptotically exact estimation approaches, such as Markov-chain Monte Carlo and full grid sampling, do not scale well to droplet-based scRNA-seq datasets that can contain thousands (and sometimes tens of thousands) of cells.

For a computationally efficient solution to estimate *P* (***λ*** | **X***, **θ***), we use variational inference, an optimization-based approximate Bayesian inference approach.

In brief, we initialize an approximate posterior distribution *q*(***λ*** *|* **X***, **θ***) that follows a transformed Beta distribution with differentiable parameters describing its shape. *q*(***λ*** *|* **X***, **θ***) is then optimized to minimize its KL divergence with *P* (***λ*** *|* **X***, **θ***) by minimizing the evidence lower bound (ELBO) objective function.

### Conversion of Bayesian ANOVA parameter posteriors to circadian cosinor parameter posteriors

Given our approximate posterior *q*(***λ*** | **X***, **θ***), for gene *j* we can generate a sampled waveform 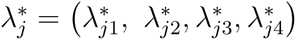, where 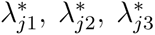, and 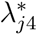 correspond to sampled gene proportions from ZT0, ZT6, ZT12, and ZT18, respectively.

Using this sampled waveform, we apply a Fast Fourier Transform, which yields sampled cosinor parameters corresponding to 12-hour and 24-hour components. We denote the sampled 24-hour circadian components for gene *j* as 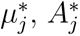, and 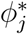, corresponding to the mesor, amplitude, and acrophase, respectively. Sampling from *q*(***λ*** *|* **X**, ***θ***) repeatedly, we can generate samples of *µ_j_, A_j_,* and *ϕ_j_* to approximate their posterior distributions.

### Identification of circadian cycling genes

Circadian cyclers were identified using three steps. First, we computed the Bayes factor of each gene using *q*(***λ*** *|* **X***, **θ***), where 5 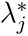 samples were drawn for each gene at each time point. Second, we computed the Bayesian evidence of each gene using an altered version of *q*(***λ*** *|* **X***, **θ***), where the 24-hour circadian component was subtracted out. To do this, 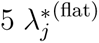 samples were drawn for each gene. Samples were then converted to the frequency domain using the Fast Fourier Transform, and the 24-hour circadian component amplitudes were set to zero. Frequency domain samples were then transformed back to the original domain using the Inverse Fast Fourier Transform, and then the Bayesian evidence was computed. Third, we compute a Bayes factor for each gene equal to the Bayesian evidence associated with *q*(***λ*** *|* **X***, **θ***) divided by the Bayesian evidence associated with the 24-hour component subtracted posterior. We refer to this Bayes factor as the circadian Bayes factor.

To further ensure that the 24-hour circadian component was the dominant frequency component, we drew 100 additional posterior waveform samples per gene, applied the Fast Fourier Transform to each, and computed the amplitude of the 24-hour component relative to other frequency components. Genes for which the 24-hour component had the largest amplitude in at least 60% of samples were retained.

Cycling genes were defined as those satisfying all of the following criteria, unless otherwise stated: (i) circadian Bayes factor *≥* 100 (log_10_ *≥* 2); (ii) 24-hour component dominant in *≥* 60% of posterior samples; (iii) detected in *>* 1% of cells and in *>* 50 cells; (iv) expected mesor *≥* 10*^−^*^6^ UMI per total library (i.e. log*e*(mesor) *≥ −*13.8).

### Gene set acrophase enrichment analysis

To identify gene sets whose members peak at coordinated times of day, we performed an acrophase enrichment analysis using two gene set databases: Reactome pathways obtained from the GSEA Molecular Signatures Database (MSigDB) and transcription factor motif target sets from the TRANSFAC and JASPAR PWM databases. Reactome pathways were filtered to exclude cancer-, disease-, and infection-related terms and restricted to sets with at most 500 member genes. For each gene set with at least 6 members among the circadian cyclers, member acrophases were binned into 12 equal-width bins spanning the 24-hour cycle (each bin = 2 hours). A likelihood ratio statistic was computed as the ratio of the observed bin density to the expected uniform density (1*/*12) at each bin, and the maximum across bins was retained as the enrichment score. Gene sets with a maximum likelihood ratio *≥* 2 were considered enriched. Enriched sets were ranked by this score and the top sets visualized as rose plots. The circular mean and circular standard deviation of member acrophases were computed for each enriched set.

### Two-group waveform enrichment analysis

To identify gene sets with sex-differential temporal expression patterns, we developed a two-group waveform enrichment analysis. For each gene, the posterior waveform was sampled independently from the two conditions (100 samples each), and the difference waveform (e.g. female *−* male) was computed sample-wise. For each gene set, the difference waveforms of member genes were averaged to obtain a gene-set-level difference waveform. To test whether the gene-set difference waveform deviates significantly from zero, we applied pointwise credible interval tests at each of the 4 time points (ZT0, ZT6, ZT12, ZT18), with Bonferroni correction for 4 comparisons. A gene set was considered significant if the 95% Bonferroni-corrected credible interval excluded zero at any time point. Significant gene sets were ranked by maximum absolute amplitude of the mean difference waveform and visualized as heatmaps.

### Intracellular clock gene phase relationship analysis

To assess whether acute misalignment altered the internal timing relationships among clock genes within SMCs (Figure 4c), we computed the relative phase delay between target clock genes and Arntl in aligned and misaligned conditions. For each condition, 30,000 posterior waveform samples were drawn for Arntl and each candidate target gene. The acrophase of each sample was estimated by applying a Fast Fourier Transform and extracting the phase of the 24-hour component. The relative phase delay was computed as the circular difference (target acrophase *−* Arntl acrophase), modulo 2*π*, yielding a posterior distribution of intracellular phase delays for each target gene in each condition.

To ensure that phase estimates were sufficiently precise for meaningful comparison, we restricted the analysis to clock genes whose posterior acrophase 95% interval was no wider than 2 hours in both aligned and misaligned conditions. The 95% interval was defined as the smallest circular arc containing 95% of the posterior acrophase samples. Changes in the relative phase delay between conditions indicate altered intracellular clock architecture beyond a simple uniform phase shift.

### Downsampling-controlled cycler count comparisons

Comparisons of the number of cycling genes between conditions can be confounded by differences in cell counts and sequencing depth, both of which affect statistical power to detect rhythmicity. To ensure fair comparisons, we applied a uniform downsampling procedure across three types of contrasts: male versus female (Figure 3b), aligned versus misaligned (Supplementary Figure 5), and wild-type versus inducible Bmal1 knockout (Supplementary Figure 11).

For each contrast, we identified the minimum number of cells at each time point across the conditions being compared within each cell type. Cells were then randomly sampled (with replacement) to this minimum at every condition-by-time-point combination, equalizing the number of cells contributing to each analysis. Next, we computed the pseudobulk UMI count (total transcripts) for each condition-by-time-point group and identified the global minimum across all groups. Total UMI counts within each group were then downsampled to this minimum via multinomial sampling, ensuring matched sequencing depth. Per-cell library sizes were recomputed from the downsampled counts. Our Bayesian non-parametric temporal regression and hypothesis testing pipeline was then rerun independently on each condition, and cycler counts were compared across a range of log_10_ circadian Bayes factor thresholds.

### Cross-sex log-likelihood analysis

To assess whether Bmal1 knockout altered the similarity of gene expression between sexes (Supplementary Figure 11c), we computed a cross-sex log-likelihood metric for each gene using the downsampled data described above (wild-type and knockout conditions matched for cell counts and library sizes across all four sex-by-genotype groups).

For each gene, 300 posterior waveform samples were drawn independently from the male wild-type, female wild-type, male knockout, and female knockout approximate posteriors. To establish a biologically meaningful scale, we first computed the male knockout effect as the difference between male knockout and male wild-type posterior waveform samples. The 95% credible interval width of this effect at each time point served as a normalization factor, ensuring that cross-sex differences were expressed relative to the magnitude of the knockout perturbation.

The cross-sex log-likelihood for each gene in the wild-type condition was computed as:

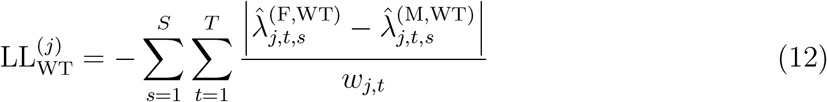

where 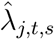 denotes the *s*-th posterior waveform sample for gene *j* at time point *t*, superscripts indicate sex and genotype, and *w_j,t_* is the 95% credible interval width of the male knockout effect at time point *t*. The knockout cross-sex log-likelihood 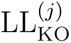 was computed analogously using female and male knockout samples. Higher (less negative) values indicate greater similarity between sexes. Genes above the *y* = *x* line have more similar expression between sexes after knockout than in wild-type, consistent with sex-specific expression being mediated through the circadian clock.

Genes were classified as wild-type cyclers (log_10_ circadian Bayes factor *≥* 10 in either male or female wild-type) or non-cyclers, and linear regression best-fit lines were computed separately for each group. Genes detected in fewer than 20% of cells in any condition, as well as hemoglobin genes, were excluded from the analysis.

### Estimating the transcript proportion – dispersion relationship

We model UMI counts as coming from a Negative Binomial (NB) data likelihood model, as it can account for the overdispersion observed in real count data. While many existing applications of NB to scRNA-seq data model dispersions in a gene-specific fashion, these estimates are known to exhibit high amounts of uncertainty. Methods such as scTransform [134] and DESeq2 [135] combat this by an empirical Bayes approach to shrink dispersion estimates, assuming genes with similar means share similar dispersions. We take this a step further and make the simplifying assumption that the transcript proportion–dispersion relationship across all genes is strictly described by a polynomial function *g**_ζ_***(*λ_ij_*) parameterized by ***ζ***. The coefficients of this polynomial, ***ζ***, are estimated as follows:

### Step 1: Estimate gene-specific proportions and dispersions

We presume gene counts are distributed according to:

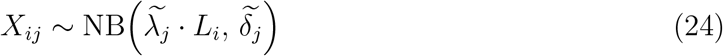

Where:

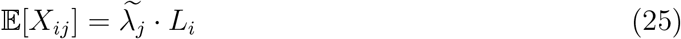

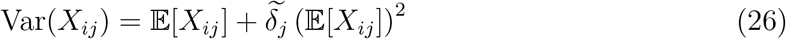

And 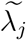 and 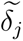 are the temporary (i.e. only used to estimate ***ζ***) gene-specific transcript proportion and dispersion for gene *j*, respectively. Under this likelihood model, for each gene we estimate the maximum likelihood transcript proportion 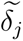 and dispersion 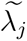.

### Step 2: Fit polynomial model describing the transcript proportion – dispersion relationship across all genes

Assume *g* is a polynomial function whose coefficients ***ζ*** defines the transcript log proportion – log dispersion relationship shared across all genes:

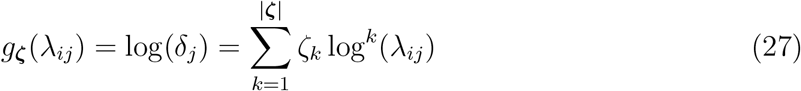

Using OLS, we estimate the coefficients ***ζ*** under the following objective:

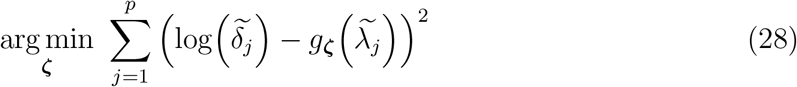

where *p* is the number of genes, and 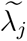 and 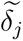 are the temporary gene-specific transcript proportion and dispersion for gene *j* estimated in Step 1.

We note that dispersion estimates may be upwardly biased as a subset of genes cycling over the circadian cycle will be fit under a mean model that assumes flat expression over the circadian cycle. However, we assume the fraction of cycling genes detected is small, and thus should not upwardly bias the dispersion estimates much.

## Acknowledgements

This work was supported by 2U54TR001878 (G.A.F), R01GM125301 (M.L.), 5TT32HL007953-22 (B.J.A.), and 2T32HG000046-21 (B.J.A.). R.L was supported by European Union’s Horizon Europe research and innovation programme under the Marie Sklodowska-Curie grant agreement No 101179447 (M1NDRHYTHM).

## Author contributions

Benjamin J. Auerbach (B.J.A.), Ronan Lordan (R.L.), Soon Yew Tang (S.Y.T), Elizabeth Hennessy (E.H), Sean T. Anderson (S.T.A.), Ujjalkumar Subhash Das (U.S.D), Ryan McConnell (R.M), Mingyao Li (M.L.), and Garret A. FitzGerald (G.A.F.) contributed to the project. The study was conceived of by G.A.F. and B.J.A. S.Y.T., R.L., R.M., E.H., U.S.D. and S.T.A. performed the mouse experiments and generated the scRNA-seq data from the mouse aorta. B.J.A. performed the data analysis, with input from M.L., R.L., S.Y.T and G.A.F. B.J.A. and R.L wrote the paper with feedback from S.Y.T., M.L. and G.A.F.

## Software availability

Scripts used to analyze the data and generate the figures can be found at https://github.com/bauerbach95/aorta_misalignment.

## Data availability

Raw sequencing data were deposited to the Sequence Read Archive under accession code SRP738214 (BioProject PRJNA1292276). The quality-controlled UMI count matrix (Ann-Data .h5ad file with cell type annotations and UMAP coordinates) and the Bayesian temporal regression outputs for each cell type and condition are available on Zenodo (https://doi.org/10.5281/zenodo.22847203). An interactive gene browser for exploring posterior waveform estimates across cell types and conditions is available at https://bauerbach95.github.io/aorta_misalignment/.

## Supplementary Figures

**Supplementary Figure 1:**
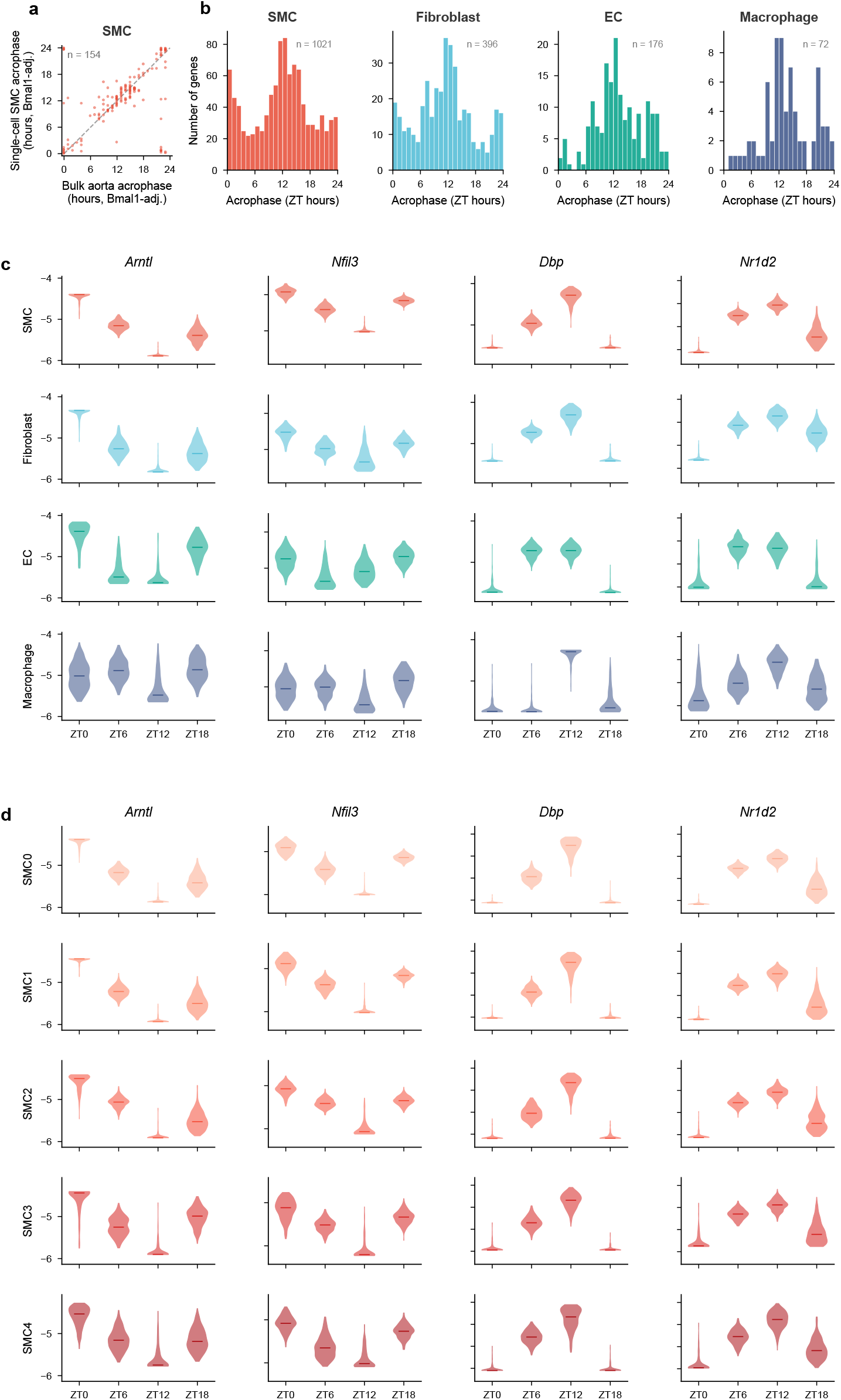
Validation of circadian cycling detection. a) Comparison of expected acrophases of SMC circadian cyclers to their peak times in bulk aorta from Zhang et al. b) Histogram of expected acrophases for cycling genes in major aortic cell types. c) Posterior waveforms of core clock genes in major aortic cell types. d) Posterior waveforms of core clock genes in SMC subtypes.

**Supplementary Figure 2:**
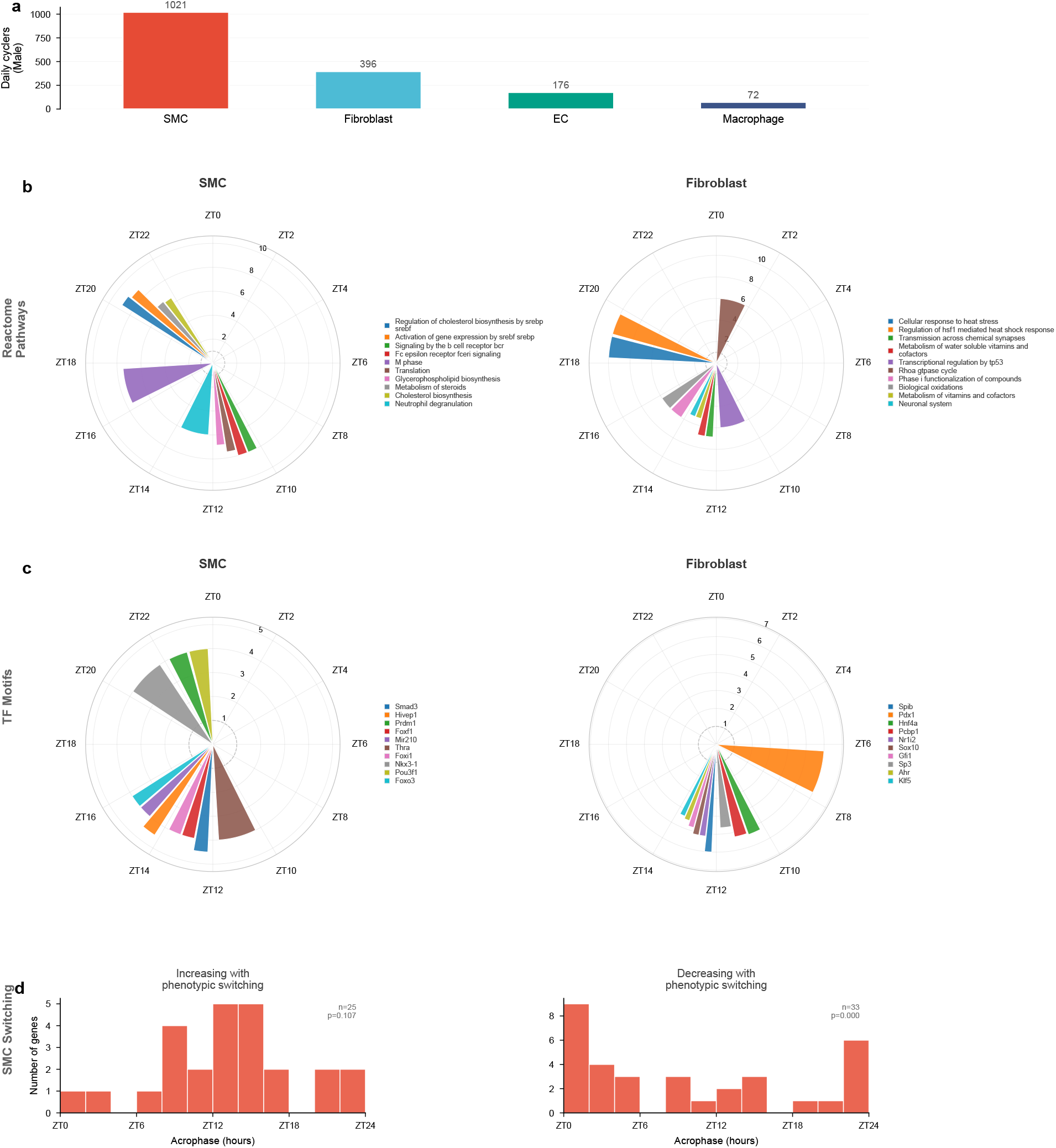
Male-specific daily cycling expression in the mouse aorta. a) Number of daily cycling genes in males for each major cell type. b) Reactome pathway enrichment among male daily cyclers in SMCs and fibroblasts. c) Transcription factor motif enrichment among male daily cyclers in SMCs and fibroblasts. d) Acrophase distributions of male SMC daily cyclers implicated in phenotypic switching, with *p*-values from window-based permutation tests.

**Supplementary Figure 3:**
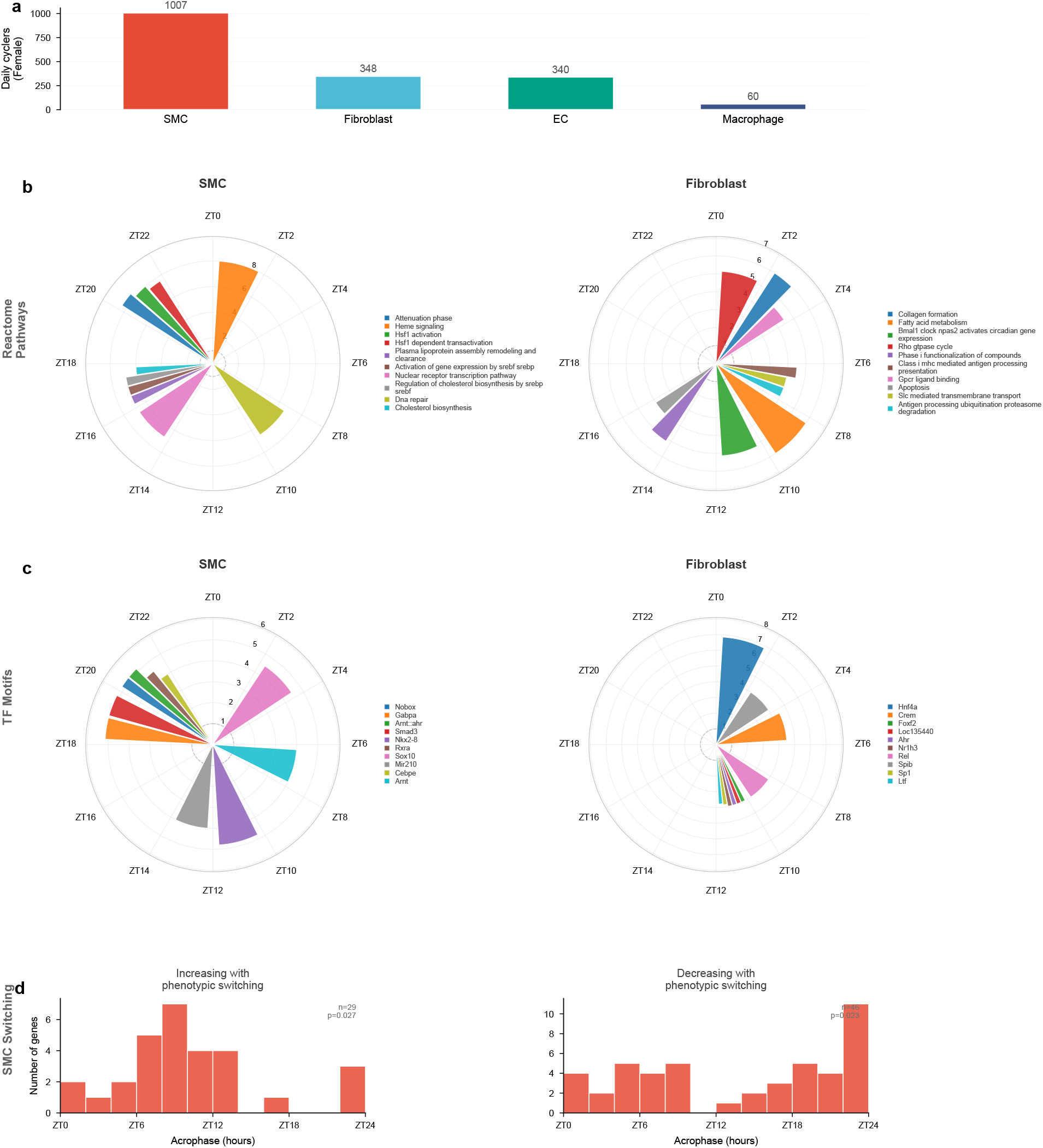
Female-specific daily cycling expression in the mouse aorta. a) Number of daily cycling genes in females for each major cell type. b) Reactome pathway enrichment among female daily cyclers in SMCs and fibroblasts. c) Transcription factor motif enrichment among female daily cyclers in SMCs and fibroblasts. d) Acrophase distributions of female SMC daily cyclers implicated in phenotypic switching, with *p*-values from window-based permutation tests.

**Supplementary Figure 4:**
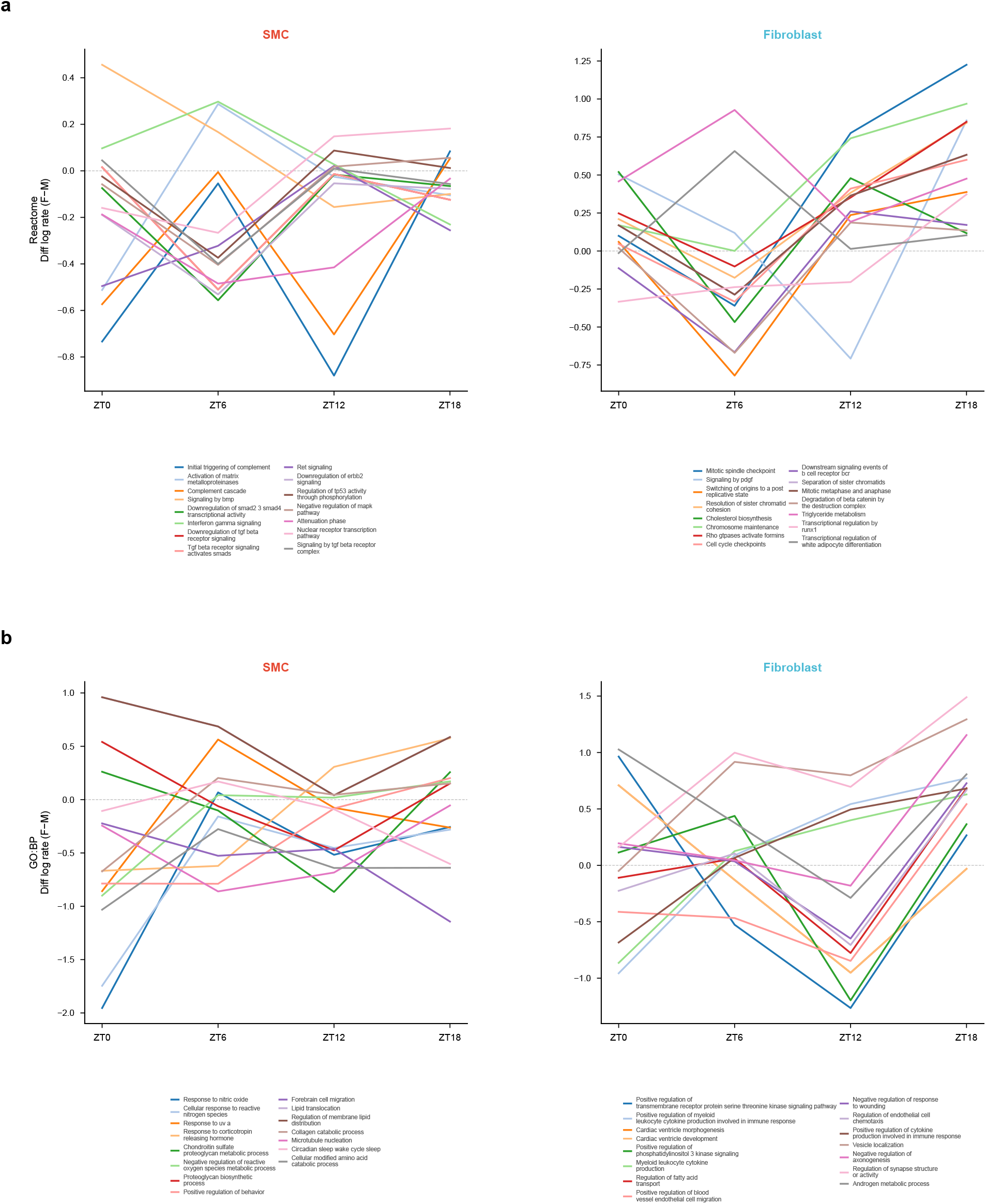
Extended waveform enrichment for sex-differential expression. Reactome pathway and GO Biological Process waveform enrichment results for sex-differential temporal expression in SMCs and fibroblasts.

**Supplementary Figure 5:**
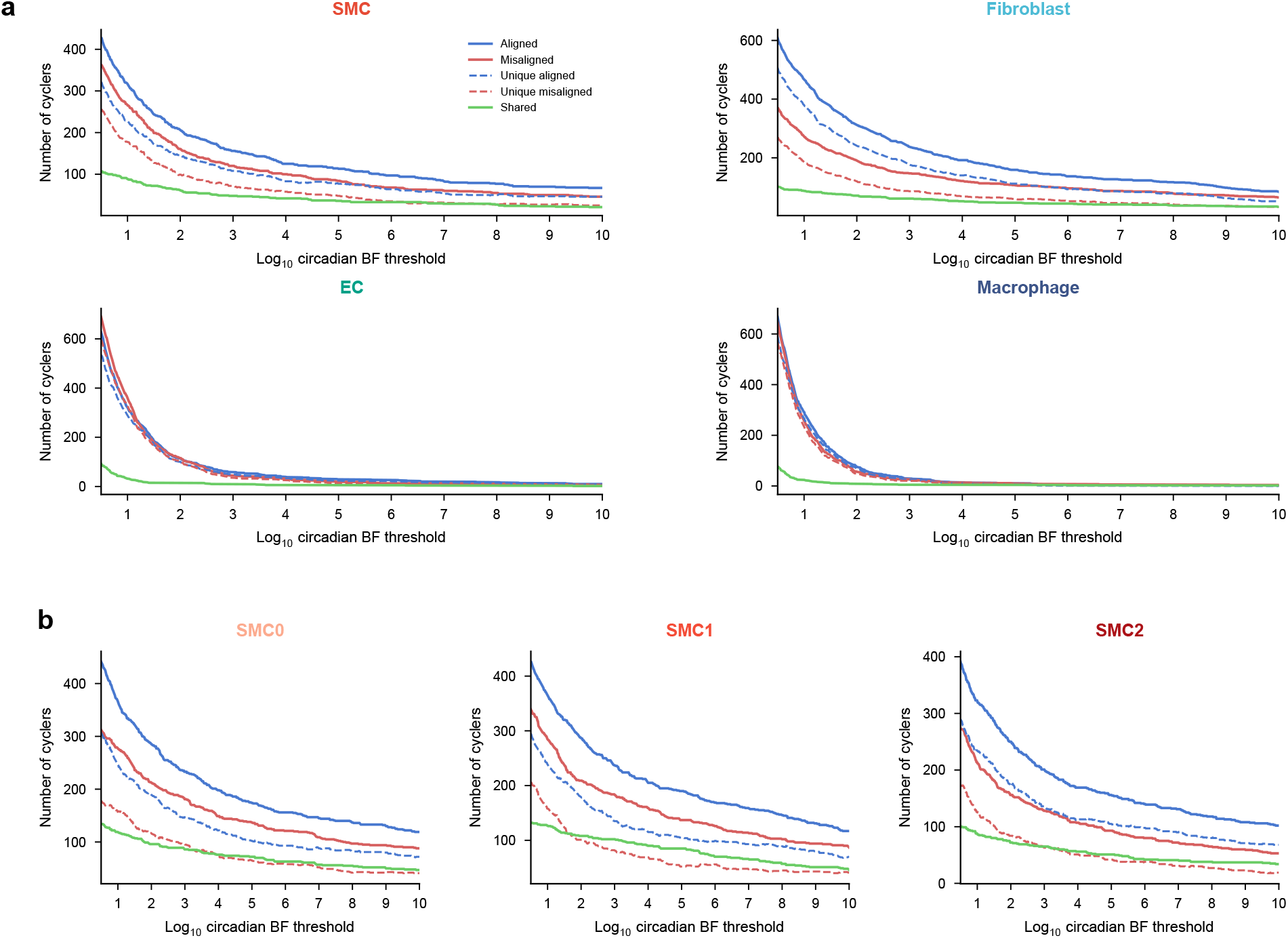
Misalignment cycling controlled for cell counts and library sizes. Within each cell type and time point, cell counts and pseudobulk UMIs were downsampled to match between aligned and misaligned male mice. a) Number of called cyclers across log_10_ circadian Bayes factor thresholds for major cell types. b) Same as a) for SMC subtypes.

**Supplementary Figure 6:**
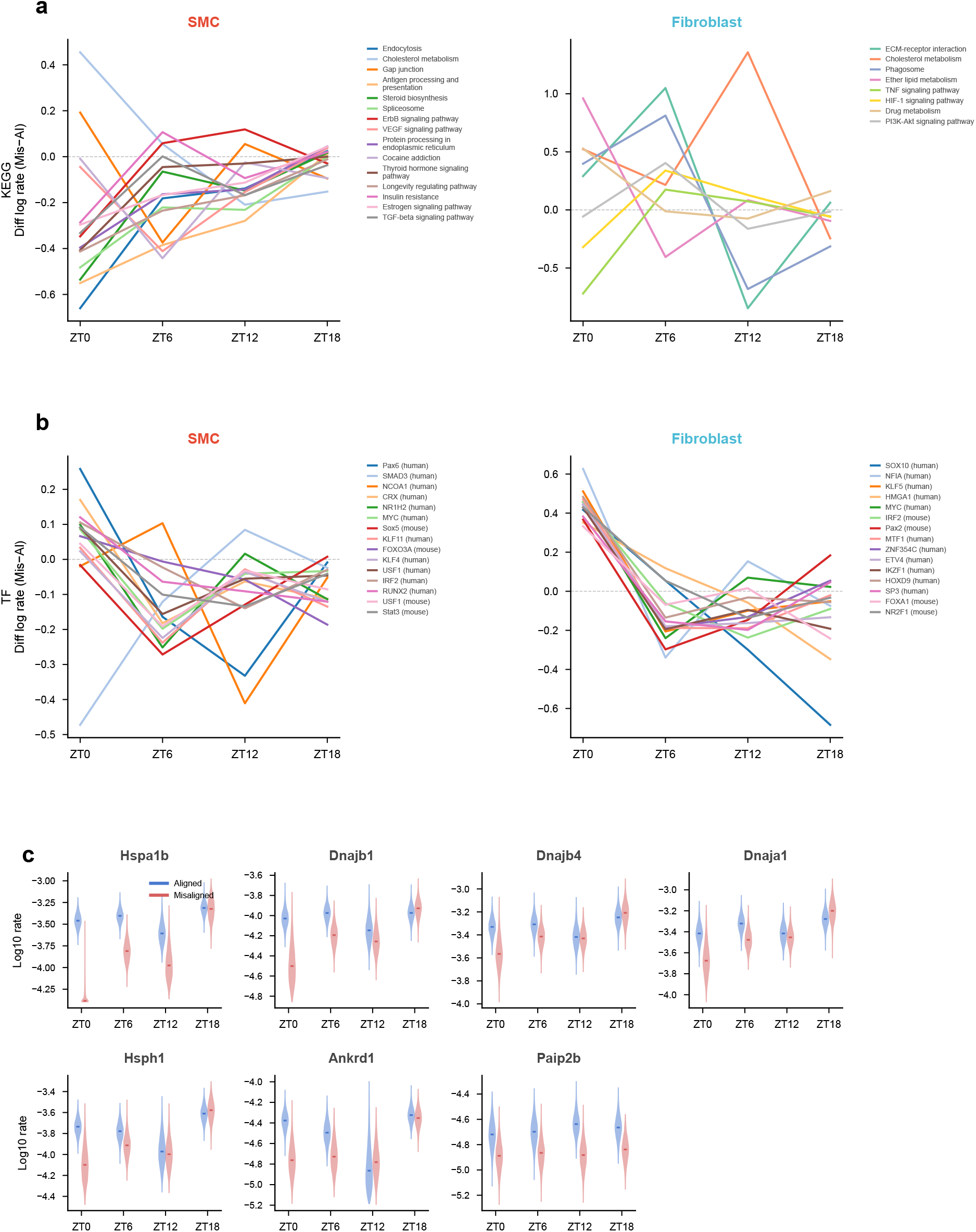
Gene set waveform enrichment and additional proteostasis gene waveforms under acute misalignment in males. a,b) Difference waveforms (misaligned minus aligned) for gene sets from a) KEGG and b) transcription factor motif databases in male SMCs and fibroblasts. Each line shows the posterior mean difference in log expression rate between misaligned and aligned conditions across zeitgeber time for a significantly enriched gene set. c) Posterior waveforms of additional proteostasis-related genes (Hspa1b, Dnajb1, Dnajb4, Dnaja1, Hsph1, Ankrd1, Paip2b) in male SMCs, comparing aligned and misaligned conditions.

**Supplementary Figure 7:**
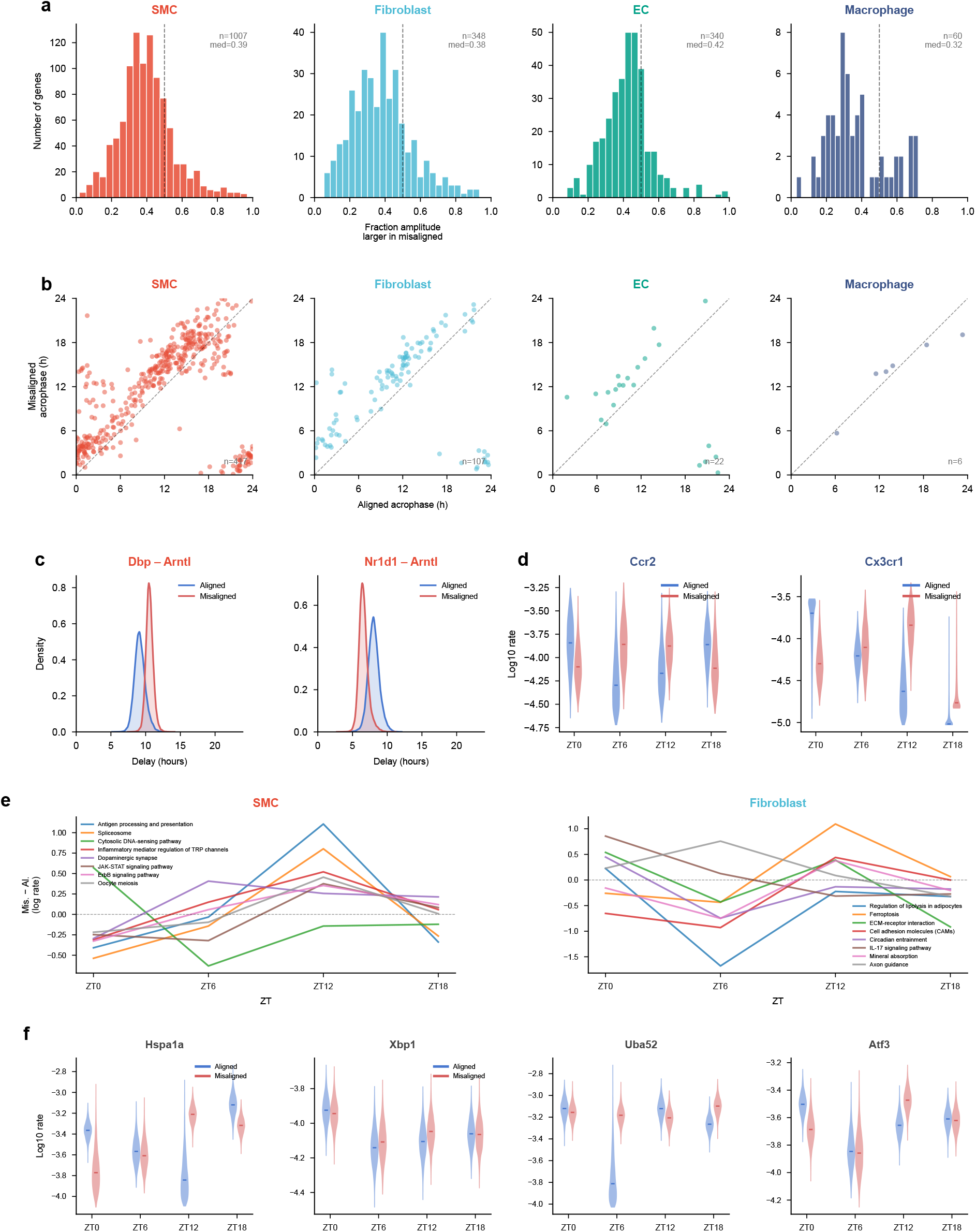
Acute misalignment effects in females. Female counterpart to Figure 4. a) Histograms of the posterior probability that misaligned amplitude exceeds aligned amplitude for each female aligned cycler, by cell type. b) Acrophase scatter plots of shared cyclers between aligned and misaligned female conditions. c) Clock gene relative timing (delay of Dbp and Nr1d1 from Arntl) in female SMCs under aligned and misaligned conditions. d) Posterior waveforms of Ccr2 and Cx3cr1 in female macrophages under aligned and misaligned conditions. e) KEGG pathway waveform enrichment (misaligned minus aligned) in female SMCs and fibroblasts. f) Posterior waveforms of key proteostasis genes (Hspa1a, Xbp1, Uba52, Atf3) in female SMCs, comparing aligned and misaligned conditions.

**Supplementary Figure 8:**
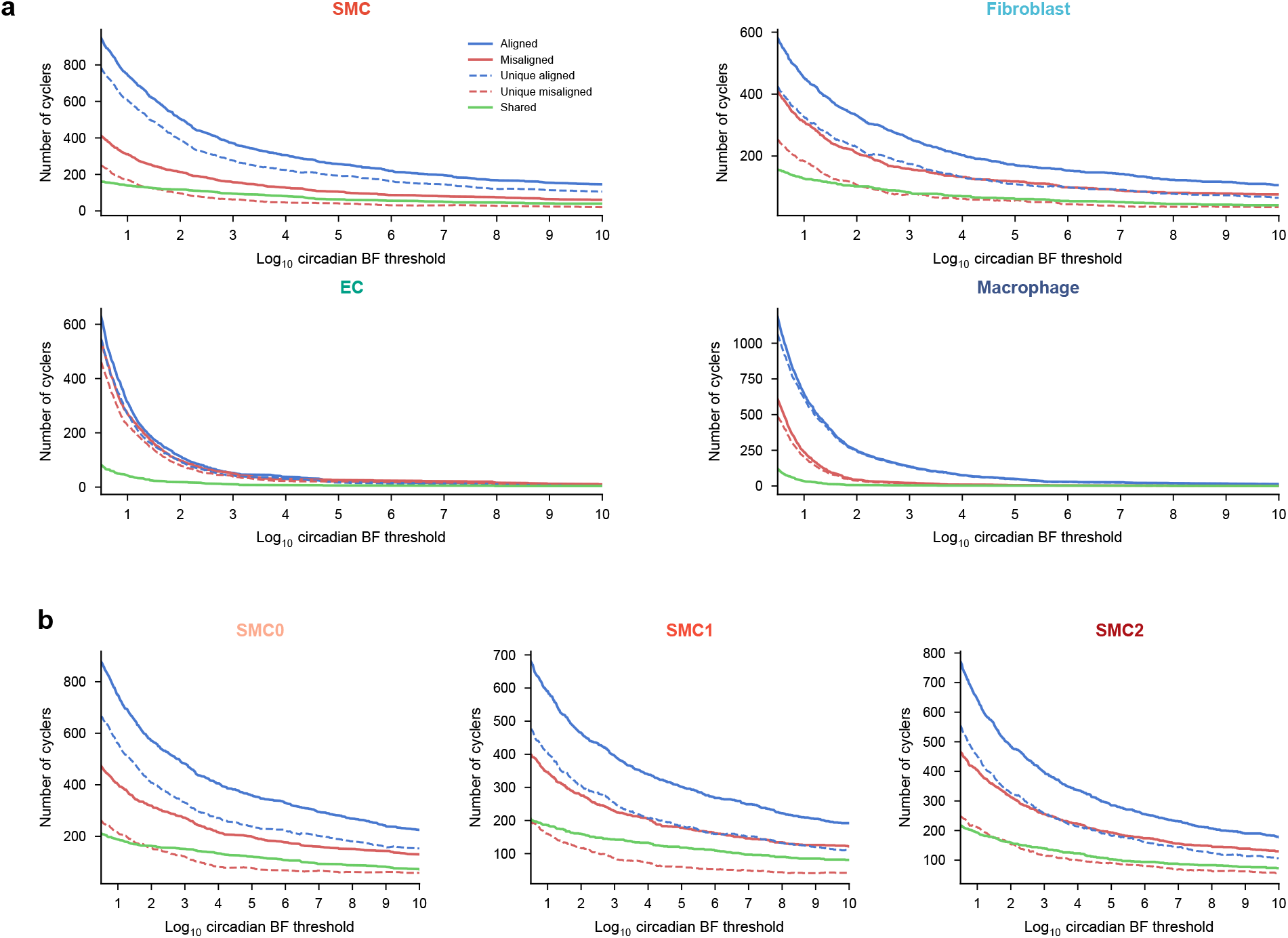
Female misalignment cycling controlled for cell counts and library sizes. Female counterpart to Supplementary Figure 5. Within each cell type and time point, cell counts and pseudobulk UMIs were downsampled to match between aligned and misaligned female mice. a) Number of called cyclers across log_10_ circadian Bayes factor thresholds for major cell types. b) Same as a) for SMC subtypes.

**Supplementary Figure 9:**
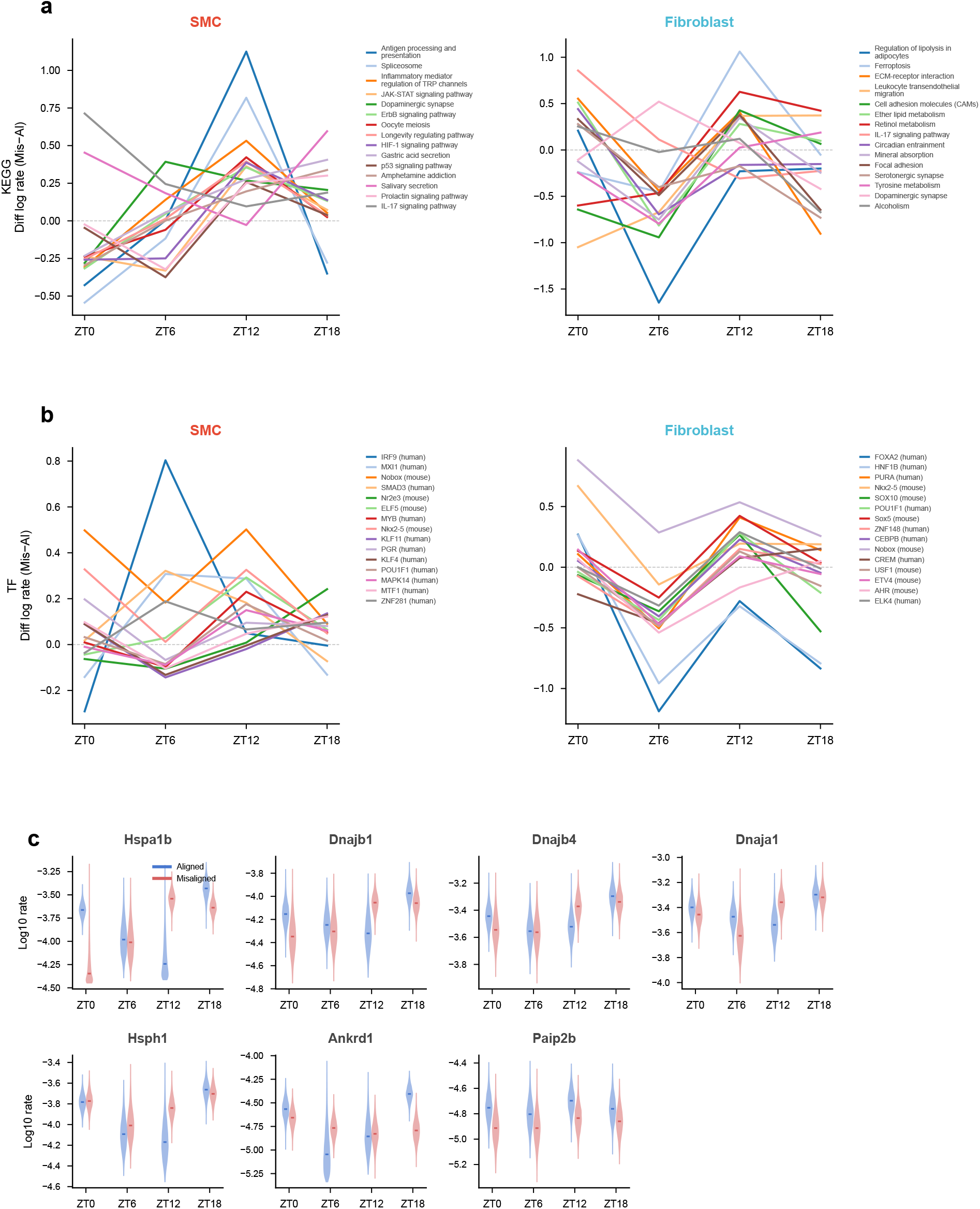
Gene set waveform enrichment and additional proteostasis gene waveforms under acute misalignment in females. Female counterpart to Supplementary Figure 6. a,b) Difference waveforms (misaligned minus aligned) for gene sets from a) KEGG and b) transcription factor motif databases in female SMCs and fibroblasts. Each line shows the posterior mean difference in log expression rate between misaligned and aligned conditions across zeitgeber time for a significantly enriched gene set. c) Posterior waveforms of additional proteostasis-related genes (Hspa1b, Dnajb1, Dnajb4, Dnaja1, Hsph1, Ankrd1, Paip2b) in female SMCs, comparing aligned and misaligned conditions.

**Supplementary Figure 10:**
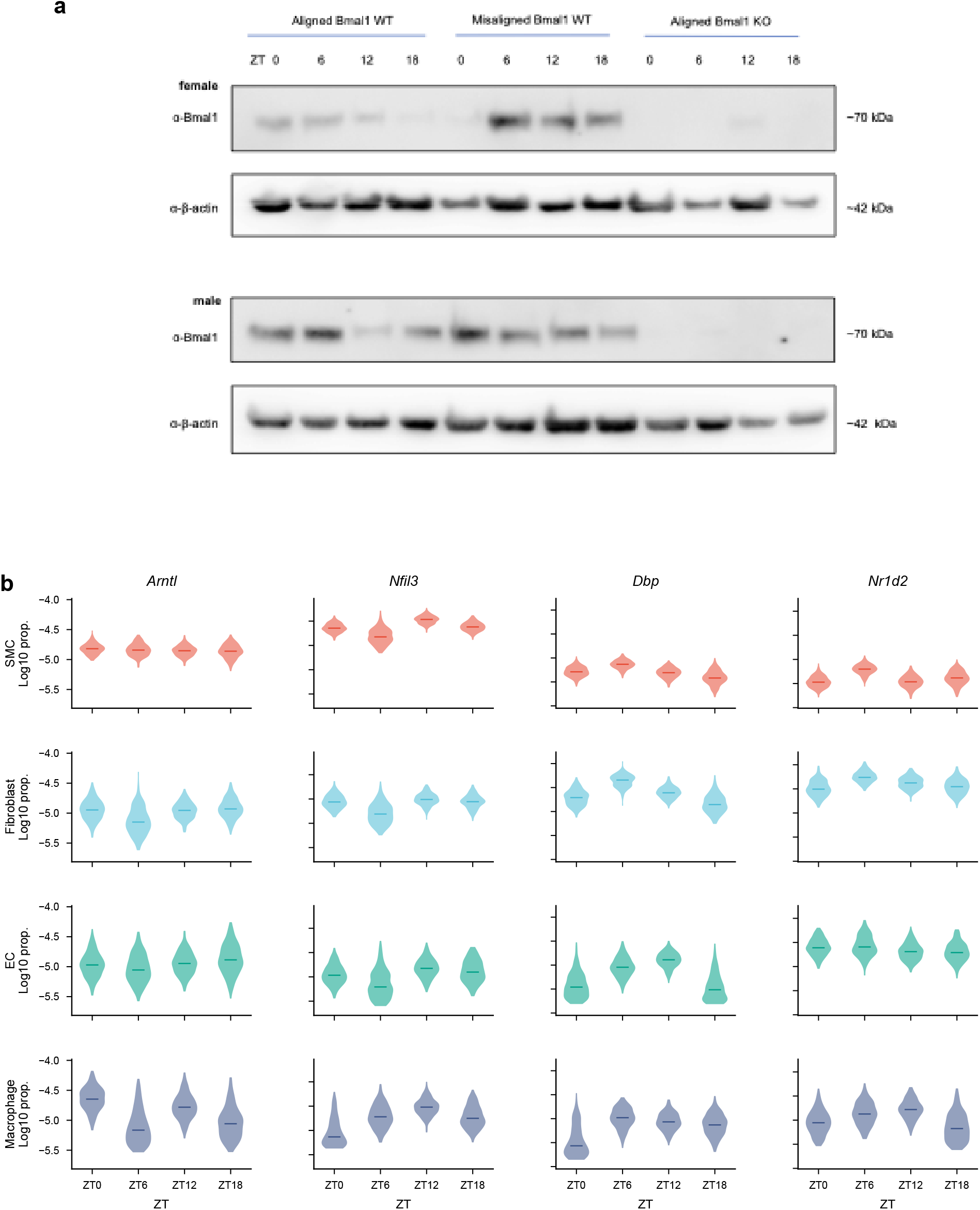
Validation of inducible Bmal1 deletion. a) Western blots of Bmal1 protein product from aorta across all conditions collected in the study. b) Posterior log_10_ proportion estimates for core circadian clock genes (Arntl, Nfil3, Dbp, Nr1d2) at each sampled time point and in each major cell type of the male Bmal1 knockout mice, showing loss of rhythmic expression.

**Supplementary Figure 11:**
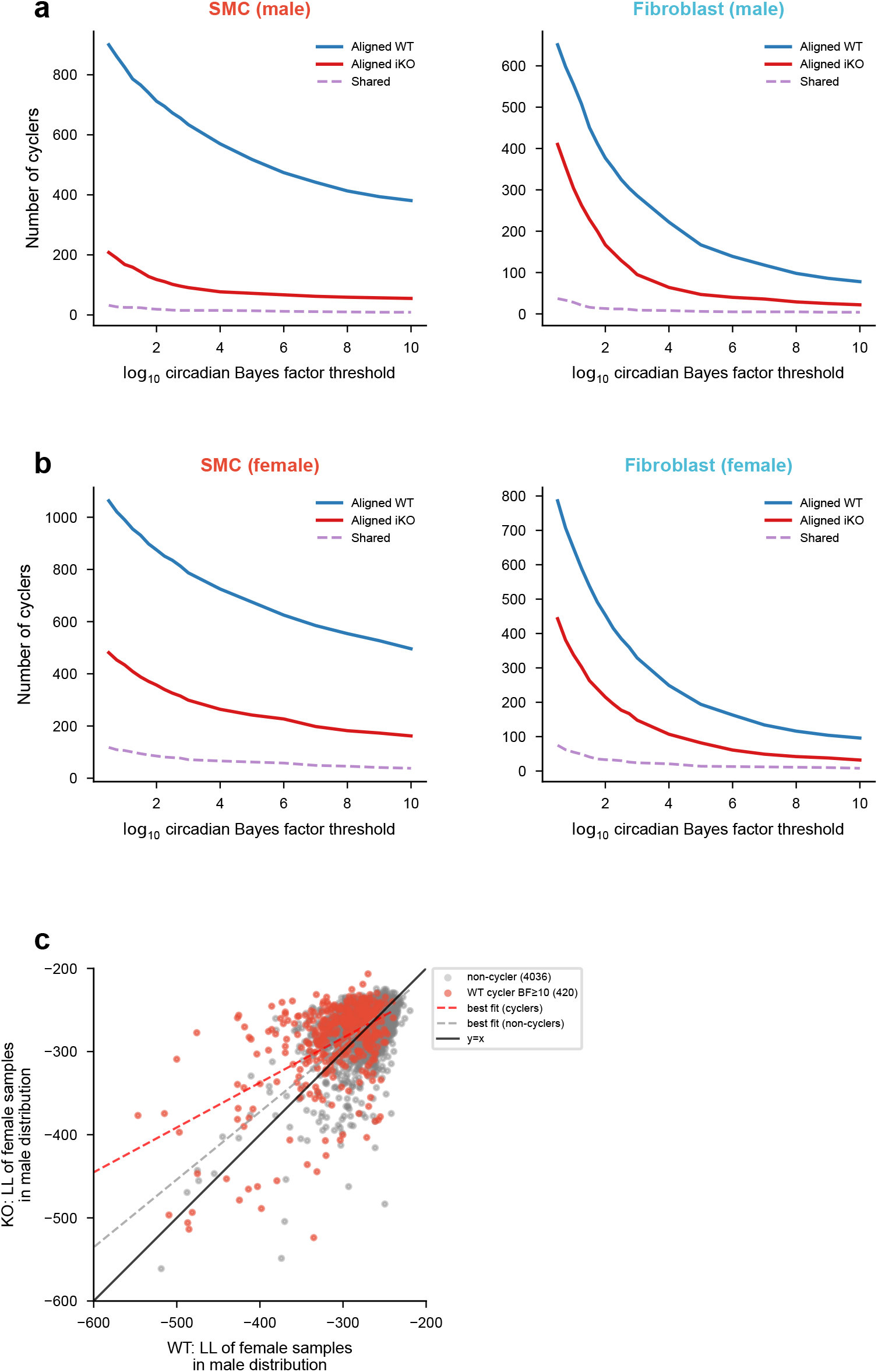
Bmal1 knockout cycling controlled for cell counts and library sizes. Within each cell type and time point, cells and pseudobulk UMI counts were downsampled to match between aligned wild-type and aligned inducible Bmal1 knockout mice. a) Number of called cyclers across log_10_ circadian Bayes factor thresholds for SMCs (left) and fibroblasts (right) in males. b) Same as (a) for females. Blue: aligned wild-type; red: aligned inducible knockout; dashed purple: shared cyclers between conditions. c) Cross-sex log-likelihood scatter for all detected SMC genes. For each gene, the similarity between female and male expression was computed in wild-type (x-axis) and knockout (y-axis) conditions by sampling from posterior waveforms and normalizing by the male knockout effect confidence interval. Points above *y* = *x* indicate more similar expression between sexes after knockout. Genes are colored by whether they are WT cyclers (red, log_10_ BF *≥* 10) or non-cyclers (grey), with separate best-fit lines for each group.

## References

[1] Panza, J. A., Epstein, S. E. & Quyyumi, A. A. Circadian Variation in Vascular Tone and Its Relation to *α*-Sympathetic Vasoconstrictor Activity. New England Journal of Medicine 325, 986–990 (1991).

[2] Bridges, A. B., McLaren, M., Saniabadi, A., Fisher, T. C. & Belch, J. J. F. Circadian variation of endothelial cell function, red blood cell deformability and dehydrothromboxane B2 in healthy volunteers. Blood Coagulation & Fibrinolysis 2, 447–452 (1991).

[3] Tunçtan, B. et al. Circadian variation of nitric oxide synthase activity in mouse tissue. Chronobiol Int 19, 393–404 (2002).

[4] Winter, C. et al. Chrono-pharmacological Targeting of the CCL2-CCR2 Axis Ameliorates Atherosclerosis. Cell Metab 28, 175–182.e5 (2018).

[5] He, W. et al. Circadian Expression of Migratory Factors Establishes Lineage-Specific Signatures that Guide the Homing of Leukocyte Subsets to Tissues. Immunity 49, 1175–1190.e7 (2018).

[6] Morris, C. J., Purvis, T. E., Hu, K. & Scheer, F. A. J. L. Circadian misalignment increases cardiovascular disease risk factors in humans. Proceedings of the National Academy of Sciences 113, E1402–E1411 (2016).

[7] Knutsson, A., Jonsson, B. G., Akerstedt, T. & Orth-Gomer, K. Increased risk of ischaemic heart disease in shift workers. The Lancet 328, 89–92 (1986).

[8] Karlsson, B., Knutsson, A. & Lindahl, B. Is there an association between shift work and having a metabolic syndrome? Results from a population based study of 27,485 people. Occup Environ Med 58, 747–752 (2001).

[9] Tüchsen, F., Hannerz, H. & Burr, H. A 12 year prospective study of circulatory disease among Danish shift workers. Occup Environ Med 63, 451–455 (2006).

[10] Lin, C. et al. Clock Gene Bmal1 Disruption in Vascular Smooth Muscle Cells Worsens Carotid Atherosclerotic Lesions. Arterioscler Thromb Vasc Biol (2022) doi:10.1161/atvbaha.121.316480.

[11] Shen, Y. et al. BMAL1 modulates smooth muscle cells phenotypic switch towards fibroblast-like cells and stabilizes atherosclerotic plaques by upregulating YAP1. Biochim Biophys Acta Mol Basis Dis 1868, (2022).

[12] Pan, X., Jiang, X. C. & Hussain, M. M. Impaired cholesterol metabolism and enhanced atherosclerosis in clock Mutant Mice. Circulation 128, 1758–1769 (2013).

[13] Cheng, B. et al. Tissue-intrinsic dysfunction of circadian clock confers transplant arteriosclerosis. Proc Natl Acad Sci U S A 108, 17147–17152 (2011).

[14] DuPont, J. J., Kenney, R. M., Patel, A. R. & Jaffe, I. Z. Sex differences in mechanisms of arterial stiffness. British Journal of Pharmacology 176, 4208–4225 (2019).

[15] Wang, M. et al. Gender heterogeneity in dyslipidemia prevalence, trends with age and associated factors in middle age rural Chinese. Lipids Health Dis 19, (2020).

[16] Palmisano, B. T., Zhu, L., Eckel, R. H. & Stafford, J. M. Sex differences in lipid and lipoprotein metabolism. Molecular Metabolism 15, 45–55 (2018).

[17] Gillis, E. E. & Sullivan, J. C. Sex Differences in Hypertension: Recent Advances. Hypertension 68, 1322–1327 (2016).

[18] Klein, S. L. & Flanagan, K. L. Sex differences in immune responses. Nature Reviews Immunology 16, 626–638 (2016).

[19] Man, J. J., Beckman, J. A. & Jaffe, I. Z. Sex as a Biological Variable in Atherosclerosis. Circ Res 1297–1319 (2020) doi:10.1161/CIRCRESAHA.120.315930.

[20] Vrijenhoek, J. E. P. et al. Sex is associated with the presence of atherosclerotic plaque hemorrhage and modifies the relation between plaque hemorrhage and cardiovascular outcome. Stroke 44, 3318–3323 (2013).

[21] Anderson, S. T. & FitzGerald, G. A. Sexual dimorphism in body clocks. Science 369, 1164–1165 (2020).

[22] Man, A. W. C., Li, H. & Xia, N. Circadian rhythm: Potential therapeutic target for atherosclerosis and thrombosis. International Journal of Molecular Sciences 22, 1–19 (2021).

[23] McAlpine, C. S. & Swirski, F. K. Circadian influence on metabolism and inflammation in atherosclerosis. Circ Res 119, 131–141 (2016).

[24] Paschos, G. K. & FitzGerald, G. A. Circadian clocks and vascular function. Circ Res 106, 833–841 (2010).

[25] Wolock, S. L., Lopez, R. & Klein, A. M. Scrublet: Computational Identification of Cell Doublets in Single-Cell Transcriptomic Data. Cell Syst 8, 281–291.e9 (2019).

[26] Lopez, R., Regier, J., Cole, M. B., Jordan, M. I. & Yosef, N. Deep generative modeling for single-cell transcriptomics. Nat Methods 15, 1053–1058 (2018).

[27] Traag, V. A., Waltman, L. & van Eck, N. J. From Louvain to Leiden: guaranteeing well-connected communities. Sci Rep 9, 1–12 (2019).

[28] Bennett, M. R., Sinha, S. & Owens, G. K. Vascular Smooth Muscle Cells in Atherosclerosis. Circ Res 118, 692–702 (2016).

[29] Trigueros-Motos, L. et al. Embryological-origin-dependent differences in homeobox expression in adult aorta: Role in regional phenotypic variability and regulation of NF-kB activity. Arterioscler Thromb Vasc Biol 33, 1248–1256 (2013).

[30] Dobnikar, L. et al. Disease-relevant transcriptional signatures identified in individual smooth muscle cells from healthy mouse vessels. Nat Commun 9, (2018).

[31] Gao, Y. K. et al. A regulator of G protein signaling 5 marked subpopulation of vascular smooth muscle cells is lost during vascular disease. PLoS One 17, (2022).

[32] Thysen, S., Cailotto, F. & Lories, R. Osteogenesis induced by frizzled-related protein (FRZB) is linked to the netrin-like domain. Laboratory Investigation 96, 570–580 (2016).

[33] Komori, T. Regulation of proliferation, differentiation and functions of osteoblasts by runx2. International Journal of Molecular Sciences 20, (2019).

[34] Lefebvre, V. & Dvir-Ginzberg, M. SOX9 and the many facets of its regulation in the chondrocyte lineage. Connective Tissue Research 58, 2–14 (2017).

[35] Pan, H. et al. Single-Cell Genomics Reveals a Novel Cell State During Smooth Muscle Cell Phenotypic Switching and Potential Therapeutic Targets for Atherosclerosis in Mouse and Human. Circulation 2060–2075 (2020) doi:10.1161/circulationaha.120.048378.

[36] Zhang, R., Lahens, N. F., Ballance, H. I., Hughes, M. E. & Hogenesch, J. B. A circadian gene expression atlas in mammals: Implications for biology and medicine. Proceedings of the National Academy of Sciences 111, 16219–16224 (2014).

[37] Wang, Y. et al. Clonally expanding smooth muscle cells promote atherosclerosis by escaping efferocytosis and activating the complement cascade. Proceedings of the National Academy of Sciences 117, 202006348 (2020).

[38] Wirka, R. C. et al. Atheroprotective roles of smooth muscle cell phenotypic modulation and the TCF21 disease gene as revealed by single-cell analysis. Nat Med (2019) doi:10.1038/s41591-019-0512-5.

[39] Yap, C., Mieremet, A., de Vries, C. J. M., Micha, D. & de Waard, V. Six shades of vascular smooth muscle cells illuminated by klf4 (Krüppel-like factor 4). Arteriosclerosis, Thrombosis, and Vascular Biology 2693–2707 (2021).

[40] Chen, E. Y., et al. Enrichr: interactive and collaborative HTML5 gene list enrichment analysis tool. http://amp.pharm.mssm.edu/Enrichr (2013).

[41] Rudic, R. D. et al. Bioinformatic analysis of circadian gene oscillation in mouse aorta. Circulation 112, 2716–2724 (2005).

[42] Millar-Craig, M., Bishop, C. & Raftery, E. B. Circadian variation of blood-pressure. The Lancet 311, 795–797 (1978).

[43] Sur, S. H., Mistlberger, R. E., Morris, M. & Morris, M. Circadian blood pressure and heart rate rhythms in mice. (1999).

[44] Scheer, F. A. J. L. et al. The human endogenous circadian system causes greatest platelet activation during the biological morning independent of behaviors. PLoS One 6, (2011).

[45] Luciano, A. K., Santana, J. M., Velazquez, H. & Sessa, W. C. Akt1 Controls the Timing and Amplitude of Vascular Circadian Gene Expression. J Biol Rhythms 32, 212–221 (2017).

[46] Adamovich, Y., Ladeuix, B., Golik, M., Koeners, M. P. & Asher, G. Rhythmic Oxygen Levels Reset Circadian Clocks through HIF1*α*. Cell Metab 25, 93–101 (2017).

[47] Manella, G. et al. Hypoxia induces a time- and tissue-specific response that elicits intertissue circadian clock misalignment. (2019) doi:10.1073/pnas.1914112117.

[48] Paroo, Z., Dipchand, E. S. & Noble, E. G. Estrogen attenuates postexercise HSP70 expression in skeletal muscle. Am J Physiol Cell Physiol 282, 245–251 (2002).

[49] Beere, H. M. et al. Heat-shock protein 70 inhibits apoptosis by preventing recruitment of procaspase-9 to the Apaf-1 apoptosome. Nature Cell Biology 2, (2000).

[50] Lu, T. S. et al. Induction of intracellular heat-shock protein 72 prevents the development of vascular smooth muscle cell calcification. Cardiovasc Res 96, 524–532 (2012).

[51] Kim, I. K., Shin, H. M. & Baek, W. Heat-shock response is associated with decreased production of interleukin-6 in murine aortic vascular smooth muscle cells. Naunyn Schmiedebergs Arch Pharmacol 371, 27–33 (2005).

[52] McCullagh, K. J. A., Cooney, R. & O’Brien, T. Endothelial nitric oxide synthase induces heat shock protein HSPA6 (HSP70B’) in human arterial smooth muscle cells. Nitric Oxide 52, 41–48 (2016).

[53] Shi, J., Yang, Y., Cheng, A., Xu, G. & He, F. Metabolism of vascular smooth muscle cells in vascular diseases. Am J Physiol Heart Circ Physiol 319, 613–631 (2020).

[54] Zheng, Y., Im, C.-N. & Seo, J.-S. Inhibitory effect of Hsp70 on angiotensin II-induced vascular smooth muscle cell hypertrophy. Exp Mol Med 38, 509–518 (2006).

[55] Guo, X. Transforming growth factor-*β* and smooth muscle differentiation. World J Biol Chem 3, 41 (2012).

[56] Ku, H. C. & Cheng, C. F. Master Regulator Activating Transcription Factor 3 (ATF3) in Metabolic Homeostasis and Cancer. Frontiers in Endocrinology 11, (2020).

[57] Yin, H.-M. et al. Activating transcription factor 3 coordinates differentiation of cardiac and hematopoietic progenitors by regulating glucose metabolism. Sci Adv 6, (2020).

[58] Miller, C. L. et al. Integrative functional genomics identifies regulatory mechanisms at coronary artery disease loci. Nat Commun 7, (2016).

[59] Wang, Y. et al. Dynamic changes in chromatin accessibility are associated with the atherogenic transitioning of vascular smooth muscle cells. Cardiovasc Res (2021) doi:10.1093/cvr/cvab347.

[60] Herring, J. A., Elison, W. S. & Tessem, J. S. Function of nr4a orphan nuclear receptors in proliferation, apoptosis and fuel utilization across tissues. Cells 8, (2019).

[61] Khambata, R. S., Panayiotou, C. M. & Hobbs, A. J. Natriuretic peptide receptor-3 underpins the disparate regulation of endothelial and vascular smooth muscle cell proliferation by C-type natriuretic peptide. Br J Pharmacol 164, 584–597 (2011).

[62] Orriols, M. et al. Down-regulation of Fibulin-5 is associated with aortic dilation: role of inflammation and epigenetics. Cardiovasc Res 110, 431–442 (2016).

[63] Spencer, J. A. et al. Altered vascular remodeling in fibulin-5-deficient mice reveals a role of fibulin-5 in smooth muscle cell proliferation and migration. Proceedings of the National Academy of Sciences 102, 2946–2951 (2005).

[64] Liu, Z. P., Wang, Z., Yanagisawa, H. & Olson, E. N. Phenotypic modulation of smooth muscle cells through interaction of Foxo4 and Myocardin. Dev Cell 9, 261–270 (2005).

[65] Hartman, R. J. G. et al. Sex-Stratified Gene Regulatory Networks Reveal Female Key-Driver Genes of Atherosclerosis Involved in Smooth Muscle Cell Phenotype Switching. Circulation 713–726 (2021) doi:10.1161/circulationaha.120.051231.

[66] Imamura, T. et al. Smad6 inhibits signalling by the TGF-*β* superfamily. Nature 389, 622–626 (1997).

[67] Miyazawa, K. & Miyazono, K. Regulation of TGF-*β* family signaling by inhibitory smads. Cold Spring Harb Perspect Biol 9, (2017).

[68] Nomiyama, T. et al. The NR4A Orphan Nuclear Receptor NOR1 Is Induced by Platelet-derived Growth Factor and Mediates Vascular Smooth Muscle Cell Proliferation. J Biol Chem 281, (2006).

[69] Janich, P. et al. The circadian molecular clock creates epidermal stem cell heterogeneity. Nature 480, 209–214 (2011).

[70] Brown, S. A. Circadian clock-mediated control of stem cell division and differentiation: Beyond night and day. Development (Cambridge) 141, 3105–3111 (2014).

[71] Zhang, Z.-B., Sinha, J., Bahrami-Nejad, Z. & Teruel, M. N. The circadian clock mediates daily bursts of cell differentiation by periodically restricting cell-differentiation commitment. Proceedings of the National Academy of Sciences 119, (2022).

[72] Yeung, C. Y. C. et al. Gremlin-2 is a BMP antagonist that is regulated by the circadian clock. Sci Rep 4, (2014).

[73] Zhang, X. et al. Sp1 Plays an Important Role in Vascular Calcification Both In Vivo and In Vitro. doi:10.1161/JAHA.117.

[74] Lyu, Q. et al. Hsp70 and NF-*κ*B mediated control of innate inflammatory responses in a canine macrophage cell line. Int J Mol Sci 21, 1–15 (2020).

[75] Cicha, I. et al. Connective tissue growth factor is overexpressed in complicated atherosclerotic plaques and induces mononuclear cell chemotaxis in vitro. Arterioscler Thromb Vasc Biol 25, 1008–1013 (2005).

[76] Trizzino, M. et al. EGR1 is a gatekeeper of inflammatory enhancers in human macrophages. Sci Adv 7, (2021).

[77] Herring, B. P., Hoggatt, A. M., Griffith, S. L., McClintick, J. N. & Gallagher, P. J. Inflammation and vascular smooth muscle cell dedifferentiation following carotid artery ligation. Physiol Genomics 49, 115–126 (2017).

[78] Balint, B. et al. Collectivization of Vascular Smooth Muscle Cells via TGF-*β*–Cadherin-11–Dependent Adhesive Switching. Arterioscler Thromb Vasc Biol 35, 1254–1264 (2015).

[79] Ribeiro-Silva, J. C., Miyakawa, A. A. & Krieger, J. E. Focal adhesion signaling: Vascular smooth muscle cell contractility beyond calcium mechanisms. Clinical Science 135, 1189–1207 (2021).

[80] He, X. et al. Activation of M3AChR (Type 3 Muscarinic Acetylcholine Receptor) and Nrf2 (Nuclear Factor Erythroid 2-Related Factor 2) Signaling by Choline Alleviates Vascular Smooth Muscle Cell Phenotypic Switching and Vascular Remodeling. Arterioscler Thromb Vasc Biol 40, 2649–2664 (2020).

[81] Johnson, P. F. Molecular stop signs: Regulation of cell-cycle arrest by C/EBP transcription factors. J Cell Sci 118, 2545–2555 (2005).

[82] Hauge, S., Macurek, L. & Syljuåsen, R. G. p21 limits S phase DNA damage caused by the Wee1 inhibitor MK1775. Cell Cycle 18, 834–847 (2019).

[83] Kavurma, M. M. & Khachigian, L. M. Sp1 inhibits proliferation and induces apoptosis in vascular smooth muscle cells by repressing p21WAF1/Cip1 transcription and cyclin D1-Cdk4-p21WAF1/Cip1 complex formation. Journal of Biological Chemistry 278, 32537–32543 (2003).

[84] Jouffe, C. et al. Disruption of the circadian clock component BMAL1 elicits an endocrine adaption impacting on insulin sensitivity and liver disease. Proceedings of the National Academy of Sciences 119, (2022).

[85] Yang, G. et al. Timing of expression of the core clock gene Bmal1 influences its effects on aging and survival. Sci Transl Med 8, 324ra16–324ra16 (2016).

[86] McGraw, A. P. et al. Aldosterone increases early atherosclerosis and promotes plaque inflammation through a placental growth factor-dependent mechanism. J Am Heart Assoc 2, (2013).

[87] Kiss, M. G. & Binder, C. J. The multifaceted impact of complement on atherosclerosis. Atherosclerosis 351, 29–40 (2022).

[88] Verdeguer, F. et al. Complement regulation in murine and human hypercholesterolemia and role in the control of macrophage and smooth muscle cell proliferation. Cardiovasc Res 76, 340–350 (2007).

[89] Lam, M. T. Y. et al. Rev-Erbs repress macrophage gene expression by inhibiting enhancer-directed transcription. Nature 498, 511–515 (2013).

[90] Apostolakis, S. & Spandidos, D. Chemokines and atherosclerosis: Focus on the CX3CL1/CX3CR1 pathway. Acta Pharmacologica Sinica 34, 1251–1256 (2013).

[91] Poupel, L. et al. Pharmacological inhibition of the chemokine receptor, CX3CR1, reduces atherosclerosis in mice. Arterioscler Thromb Vasc Biol 33, 2297–2305 (2013).

[92] Timmons, G. A., O’Siorain, J. R., Kennedy, O. D., Curtis, A. M. & Early, J. O. Innate Rhythms: Clocks at the Center of Monocyte and Macrophage Function. Frontiers in Immunology 11, 1743 (2020).

[93] Yasumoto, H. et al. Dominant negative c-Jun gene transfer inhibits vascular smooth muscle cell proliferation and neointimal hyperplasia in rats. Gene Therapy 8, (2001).

[94] Khachigian, L. M., Fahmy, R. G., Zhang, G., Bobryshev, Y. V. & Kaniaros, A. c-Jun regulates vascular smooth muscle cell growth and neointima formation after arterial injury. Journal of Biological Chemistry 277, 22985–22991 (2002).

[95] Das, S. et al. Regulation of angiotensin II actions by enhancers and super-enhancers in vascular smooth muscle cells. Nat Commun 8, (2017).

[96] Thompson, M. R., Xu, D. & Williams, B. R. G. ATF3 transcription factor and its emerging roles in immunity and cancer. Journal of Molecular Medicine 87, 1053–1060 (2009).

[97] Herbert, J. M., Lamarche, I., Prabonnaud, V., Dol, F. & Gauthier, T. Tissue-type plasminogen activator is a potent mitogen for human aortic smooth muscle cells. Journal of Biological Chemistry 269, 3076–3080 (1994).

[98] Ishigami, M., Swertfeger, D. K., Hui, M. S., Granholm, N. A. & Hui, D. Y. Apolipoprotein E Inhibition of Vascular Smooth Muscle Cell Proliferation but Not the Inhibition of Migration Is Mediated Through Activation of Inducible Nitric Oxide Synthase. Arterioscler Thromb Vasc Biol (2000).

[99] Lee, J. H. et al. Mig-6 gene knockout induces neointimal hyperplasia in the vascular smooth muscle cell. Dis Markers 2014, (2014).

[100] Demirel, E. et al. Rgs5 attenuates baseline activity of erk1/2 and promotes growth arrest of vascular smooth muscle cells. Cells 10, (2021).

[101] Daniel, J.-M. et al. Regulator of G-Protein Signaling 5 Prevents Smooth Muscle Cell Proliferation and Attenuates Neointima Formation. Arterioscler Thromb Vasc Biol 36, 317–327 (2016).

[102] Ji, Y. et al. Pharmacological Targeting of Plasminogen Activator Inhibitor-1 Decreases Vascular Smooth Muscle Cell Migration and Neointima Formation. Arterioscler Thromb Vasc Biol 36, 2167–2175 (2016).

[103] Tong, Y. et al. Exosome-mediated transfer of ACE (angiotensin-converting enzyme) from adventitial fibroblasts of spontaneously hypertensive rats promotes vascular smooth muscle cell migration. Hypertension 72, 881–888 (2018).

[104] Huang, C., Santofimia-Castaño, P. & Iovanna, J. NUPR1: A critical regulator of the antioxidant system. Cancers 13, (2021).

[105] Serrano, R. L., Yu, W., Graham, R. M., Liu-Bryan, R. & Terkeltaub, R. A vascular smooth muscle cell X-box binding protein 1 and transglutaminase 2 regulatory circuit limits neointimal hyperplasia. PLoS One 14, (2019).

[106] Takaguri, A., Kubo, T., Mori, M. & Satoh, K. The protective role of YAP1 on ER stress-induced cell death in vascular smooth muscle cells. Eur J Pharmacol 815, 470– 477 (2017).

[107] Berlanga, J. J., Baass, A. & Sonenberg, N. Regulation of poly(A) binding protein function in translation: Characterization of the Paip2 homolog, Paip2B. RNA 12, 1556–1568 (2006).

[108] Lecker, S. H., Goldberg, A. L. & Mitch, W. E. Protein degradation by the ubiquitin-proteasome pathway in normal and disease states. Journal of the American Society of Nephrology 17, 1807–1819 (2006).

[109] Kobayashi, M. et al. The ubiquitin hybrid gene UBA52 regulates ubiquitination of ribosome and sustains embryonic development. Sci Rep 6, (2016).

[110] Zeng, M. et al. CRL4Wdr70 regulates H2B monoubiquitination and facilitates Exo1dependent resection. Nat Commun 7, (2016).

[111] Zeng, M., Tang, Z., Guo, L., Wang, X. & Liu, C. Wdr70 regulates histone modification and genomic maintenance in fission yeast. Biochim Biophys Acta Mol Cell Res 1867, (2020).

[112] Baeten, J. T. & Lilly, B. Differential regulation of NOTCH2 and NOTCH3 contribute to their unique functions in vascular smooth muscle cells. Journal of Biological Chemistry 290, 16226–16237 (2015).

[113] Boucher, J. M., Harrington, A., Rostama, B., Lindner, V. & Liaw, L. A receptor-specific function for Notch2 in mediating vascular smooth muscle cell growth arrest through cyclin-dependent kinase inhibitor 1B. Circ Res 113, 975–985 (2013).

[114] Karakaya, C. et al. Notch signaling regulates strain-mediated phenotypic switching of vascular smooth muscle cells. Front Cell Dev Biol 10, (2022).

[115] Schultz, K., Murthy, V., Tatro, J. B. & Beasley, D. Prolyl hydroxylase 2 deficiency limits proliferation of vascular smooth muscle cells by hypoxia-inducible factor-1-dependent mechanisms. Am J Physiol Lung Cell Mol Physiol 296, 921–927 (2009).

[116] Zhao, Y. et al. Rho-associated protein kinase isoforms stimulate proliferation of vascular smooth muscle cells through ERK and induction of cyclin D1 and PCNA. Biochem Biophys Res Commun 432, 488–493 (2013).

[117] Matsumae, H. et al. CCN1 knockdown suppresses neointimal hyperplasia in a rat artery balloon injury model. Arterioscler Thromb Vasc Biol 28, 1077–1083 (2008).

[118] Forrest, S. & McNamara, C. Id family of transcription factors and vascular lesion formation. Arteriosclerosis, Thrombosis, and Vascular Biology 24, 2014–2020 (2004).

[119] Forrest, S. T., Taylor, A. M., Sarembock, I. J., Perlegas, D. & McNamara, C. A. Phosphorylation regulates Id3 function in vascular smooth muscle cells. Circ Res 95, 557–559 (2004).

[120] Lowery, J. W. et al. ID family protein expression and regulation in hypoxic pulmonary hypertension. Am J Physiol Regul Integr Comp Physiol 299, 1463–1477 (2010).

[121] Tejera-Muñoz, A. et al. Ccn2 increases tgf-*β* receptor type ii expression in vascular smooth muscle cells: Essential role of ccn2 in the tgf-*β* pathway regulation. Int J Mol Sci 23, (2022).

[122] Chen, C. C. & Lau, L. F. Functions and mechanisms of action of CCN matricellular proteins. International Journal of Biochemistry and Cell Biology 41, 771–783 (2009).

[123] Suehiro, J. I., Hamakubo, T., Kodama, T., Aird, W. C. & Minami, T. Vascular endothelial growth factor activation of endothelial cells is mediated by early growth response-3. Blood 115, 2520–2532 (2010).

[124] Liu, D., Evans, I., Britton, G. & Zachary, I. The zinc-finger transcription factor, early growth response 3, mediates VEGF-induced angiogenesis. Oncogene 27, 2989–2998 (2008).

[125] Phng, L. K. et al. Nrarp Coordinates Endothelial Notch and Wnt Signaling to Control Vessel Density in Angiogenesis. Dev Cell 16, 70–82 (2009).

[126] Harjes, U., Bridges, E., McIntyre, A., Fielding, B. A. & Harris, A. L. Fatty acid-binding protein 4, a point of convergence for angiogenic and metabolic signaling pathways in endothelial cells. Journal of Biological Chemistry 289, 23168–23176 (2014).

[127] Harjes, U. et al. Antiangiogenic and tumour inhibitory effects of downregulating tumour endothelial FABP4. Oncogene 36, 912–921 (2017).

[128] Kanemaru, H. et al. BATF2 activates DUSP2 gene expression and up-regulates NF-*κ*B activity via phospho-STAT3 dephosphorylation. Int Immunol 30, 255–265 (2018).

[129] Jeffrey, K. L. et al. Positive regulation of immune cell function and inflammatory responses by phosphatase PAC-1. Nat Immunol 7, 274–283 (2006).

[130] Rigo, A. et al. Macrophages may promote cancer growth via a GM-CSF/HB-EGF paracrine loop that is enhanced by CXCL12. Mol Cancer 9, (2010).

[131] Edwards, J. P., Zhang, X. & Mosser, D. M. The Expression of Heparin-Binding Epidermal Growth Factor-Like Growth Factor by Regulatory Macrophages. The Journal of Immunology 182, 1929–1939 (2009).

[132] Higashiyama, S., Abraham, J. A., Miller, J., Fiddes, J. C. & Klagsbrun, M. A Heparin-Binding Growth Factor Secreted by Macrophage-Like Cells That Is Related to EGF. Science 251, 936–939 (1991).

[133] Xia, C., Braunstein, Z., Toomey, A. C., Zhong, J. & Rao, X. S100 proteins as an important regulator of macrophage inflammation. Frontiers in Immunology 8, (2018).

[134] Hafemeister, C. & Satija, R. Normalization and variance stabilization of single-cell RNA-seq data using regularized negative binomial regression. Genome Biol 20, 1–15 (2019).

[135] Love, M. I., Huber, W. & Anders, S. Moderated estimation of fold change and dispersion for RNA-seq data with DESeq2. Genome Biol 15, 1–21 (2014).

[136] Hatakeyama, T., Pappas, P. J., Hobson, R. W., Boric, M. P., Sessa, W. C. & Duran, W. N. Endothelial nitric oxide synthase regulates microvascular hyperpermeability in vivo. J Physiol 574, 275–281 (2006).

[137] Cyr, A. R., Huckaby, L. V., Shiva, S. S. & Zuckerbraun, B. S. Nitric Oxide and Endothelial Dysfunction. Crit Care Clin 36, 307–321 (2020).

[138] Anderson, S. T. et al. Sexual dimorphism in the response to chronic circadian misalignment on a high-fat diet. Sci Transl Med 15, eabo2022 (2023).

[139] Paschos, G. K., Lordan, R. & FitzGerald, G. A. Intersection of sex and circadian biology. Curr Opin Physiol 45, 100834 (2025).

[140] Tang, S. Y., et al. Differential Impact In Vivo of Pf4-Cre-Mediated and Gp1ba-Cre-Mediated Depletion of Cyclooxygenase-1 in Platelets in Mice. Arterioscler Thromb Vasc Biol 44, 1393–1406 (2024).

[141] Yu, Y. et al. Vascular COX-2 Modulates Blood Pressure and Thrombosis in Mice. Sci Transl Med 4, 132ra54 (2012).

[142] Tang, S. Y. et al. Cardiovascular Consequences of Prostanoid I Receptor Deletion in Microsomal Prostaglandin E Synthase-1-Deficient Hyperlipidemic Mice. Circulation 134, 328–338 (2016).

[143] Wick, M. J., Harral, J. W., Loomis, Z. L. & Dempsey, E. C. An Optimized Evans Blue Protocol to Assess Vascular Leak in the Mouse. J Vis Exp (139), e57037 (2018).

[144] LeGates, T. A., Dunn, D. & Weber, E. T. Accelerated re-entrainment to advanced light cycles in BALB/cJ mice. Physiol. Behav. 98, 427–432 (2009).

[145] Davidson, A. J. et al. Visualizing jet lag in the mouse suprachiasmatic nucleus and peripheral circadian timing system. Eur. J. Neurosci. 29, 171–180 (2009).

[146] Yamazaki, S. et al. Resetting central and peripheral circadian oscillators in transgenic rats. Science 288, 682–685 (2000).

